# Dynamics of amylopectin granule accumulation during the course of the chronic *Toxoplasma* infection is linked to intra-cyst bradyzoite replication

**DOI:** 10.1101/2024.09.02.610794

**Authors:** Aashutosh Tripathi, Ryan W. Donkin, Joy S. Miracle, Robert D. Murphy, Matthew S. Gentry, Abhijit Patwardhan, Anthony P. Sinai

## Abstract

The contribution of amylopectin granules (AG), comprised of a branched chain storage homopolymer of glucose, to the maintenance and progression of the chronic *Toxoplasma gondii* infection has remained undefined. Here we describe the role of AG in the physiology of encysted bradyzoites by using a custom developed imaging-based application AmyloQuant that permitted quantification of relative levels of AG within *in vivo* derived tissue cysts during the initiation and maturation of the chronic infection. Our findings establish that AG are dynamic entities, exhibiting considerable heterogeneity among tissue cysts at all post infection time points examined. Quantification of relative AG levels within tissue cysts exposes a previously unrecognized temporal cycle defined by distinct phases of AG accumulation and utilization over the first 6 weeks of the chronic phase. This AG cycle is temporally coordinated with overall bradyzoite mitochondrial activity implicating amylopectin in the maintenance and progression of the chronic infection. In addition, the staging of AG accumulation and its rapid utilization within encysted bradyzoites was associated with a burst of coordinated replication. As such our findings suggest that AG levels within individual bradyzoites, and across bradyzoites within tissue cysts may represent a key component in the licensing of bradyzoite replication, intimately linking stored metabolic potential to the course of the chronic infection. This extends the impact of AG beyond the previously assigned role that focused exclusively on parasite transmission. These findings force a fundamental reassessment of the chronic *Toxoplasma* infection, highlighting the critical need to address the temporal progression of this crucial stage in the parasite life cycle.

## Introduction

The broad host range and persistence of the protozoan parasite *Toxoplasma gondii* contribute to its success as a pathogen (1, 2). The ability of *Toxoplasma gondii* to form tissue cysts in its asexual cycle is central to its long term persistence (3, 4). Tissue cysts serve as a vehicle for transmission due to carnivory, with the option of entering the sexual cycle when the carnivore is a felid (4). Tissue cysts contain several dozen to hundreds of genetically clonal organisms termed bradyzoites (5, 6). Long considered to be dormant, our work established that encysted bradyzoites retain considerable metabolic potential including the ability to replicate by endodyogeny (3, 5).

Among the morphological features associated with encysted bradyzoites is the presence of electron lucent cytoplasmic inclusions (7) that have subsequently been identified as amylopectin granules (AG) (8–10). These inclusions, which are restricted to transmission forms of the parasite, the bradyzoite and sporozoite, were noted to have their evolutionary origins with the secondary endosymbiotic event associated with the acquisition of the apicoplast, a relict genome containing plastid derived for the ancestor of red alga (9, 11). In fact, the evolution of glycogen and starch metabolism in eukaryotes provides molecular clues to the basis of plastid endosymbiosis, linking them as is the case in Apicomplexa (12). The absence of AG within tachyzoites, as well as the erroneous view that bradyzoites were replication incompetent, cemented the view that AG did not play any role during the chronic infection (8, 13). Rather, AG serves as a ready source of energy and metabolic potential to promote transmission to a new host and/or reactivation events within the original host both of which are associated with conversion to tachyzoites (8, 13). Our finding that bradyzoites are able to replicate within tissue cysts (5) prompted us to examine whether AG played any role in the progression of the chronic infection impacting energy metabolism and replication potential.

Early morphological data established that the levels and distribution of AG within encysted bradyzoites is not uniform (7, 14, 15). To capture and quantify the relative levels of AG within encysted bradyzoites we developed AmyloQuant, an imaging-based application to establish the levels and distribution of AG within tissue cysts following optimized labeling with Periodic Acid Schiff (PAS) reagent (16). Quantification of relative AG levels using AmyloQuant exposed a remarkable level of diversity in AG across tissue cysts at all time points tested. The data generated reveal that AG is dynamic, with patterns defining previously unrecognized phases in the progression of the chronic infection within the infected murine brain. Furthermore, our data establish that as a polymer of glucose, patterns for AG accumulation and breakdown correlate with levels of bradyzoite mitochondrial activity and the capacity for both sporadic and coordinated intra-cyst replication. These data suggest that such events are not random, but rather are responsive to a potentially preprogrammed temporal cycle associated with the progression of the chronic infection. Together these findings greatly expand our understanding of the contribution of AG, revealing their role in the progression and maintenance of the chronic infection by serving as a potential energy and metabolic resource to license demanding processes, such as replication. We further find that the presence of a potential temporal AG cycle reveals patterns in what otherwise appear as heterogenous populations governing overall energetic, metabolic and replicative potential of the bradyzoites within tissue cysts. As amylopectin dynamics emerge to play an important role in the normal progression of the chronic infection, perturbation of this pathway may present an opportunity for therapeutic intervention (17).

## Results

### Amylopectin levels vary within encysted bradyzoites in vivo

Amylopectin granules (AG) were first recognized as electron lucent cytoplasmic inclusions within encysted bradyzoites *in vivo* (7). AG is comprised of a branched homopolymer of glucose arranged as repeating α1,4 linked chains interconnected with α1,6 branch points (**Fig. 1A**) (10). AG is similar to plant starch and as such the parasite encodes all the genes needed for AG/starch synthesis (9, 18). AG synthesis is initiated with glucose 6-P being converted to glucose 1 phosphate by phosphoglucomutases (PGM1/2) (19, 20) which is activated with a UDP or ADP forming the nucleotide sugar (21) that is the substrate for the starch/ glycogen synthase (8, 22). As the linear polymer and branched chains are synthesized, chain winding results in the expulsion of water, generating a layer of insoluble starch as the granule grows (18) (**Fig. 1B**). Mobilization of glucose from AG necessitates the unwinding of the glucan chain, allowing access to amylases (23). This is facilitated by a glucan water dikinase (TgGWD), a unique enzyme that transfers the β-phosphate from ATP to the glucan chain (21, 24, 25). As amylase progressively releases glucose, their progress is inhibited by the presence of the phosphorylated glucose, necessitating the activity of a glucan phosphatase TgLaforin (17, 26, 27), establishing a glucan phosphorylation-dephosphorylation cycle which, in conjunction with the amylases, releases stored glucose for downstream functions.

**Figure 1.**
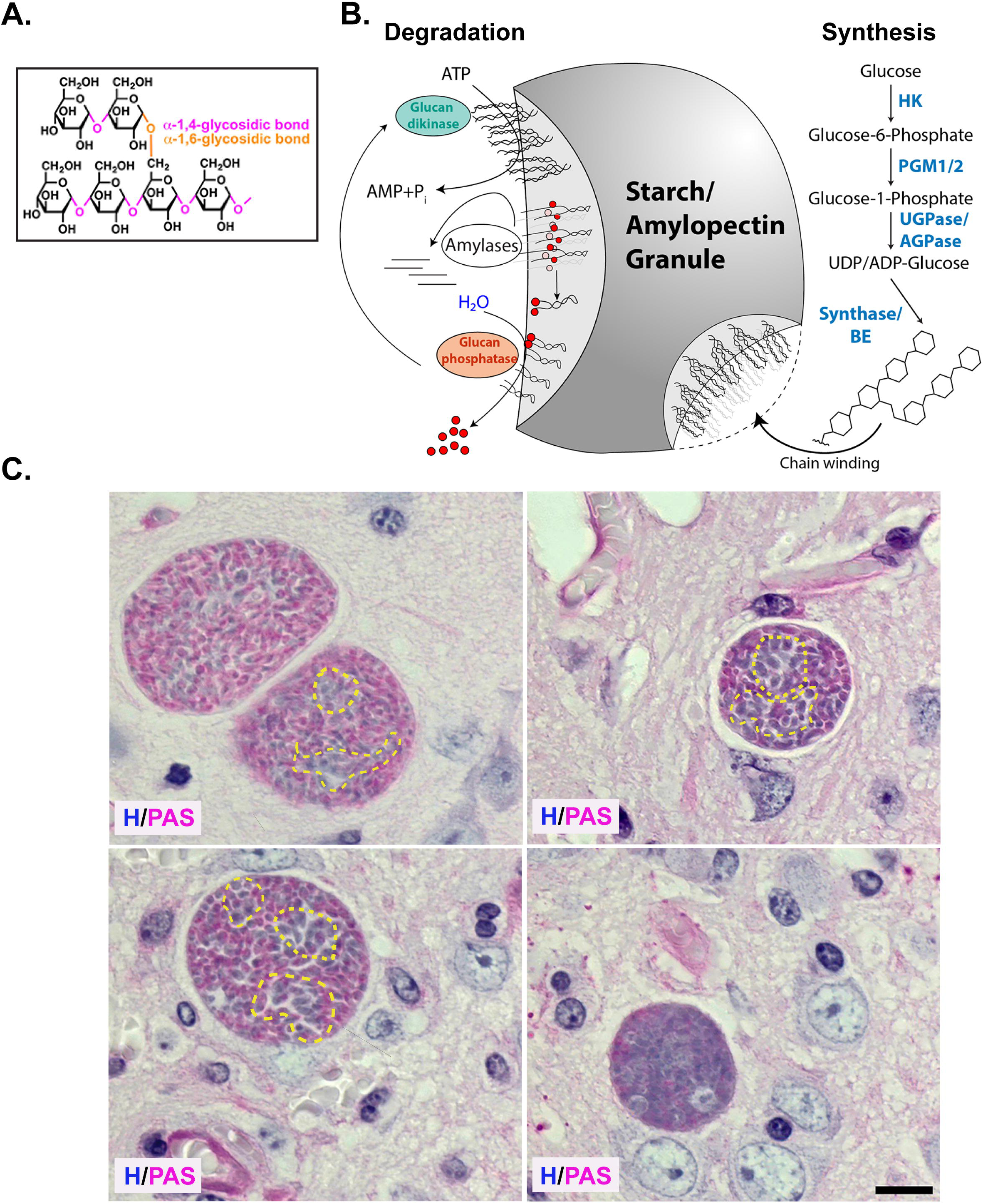
Heterogenous distribution of amylopectin granules within encysted bradyzoites in vivo. (**A**) Amylopectin is an α1,4-linked linear glucose polymer connected with α1,6 branched linkages. (Β) Steady state levels of amylopectin in water insoluble stach/amylopectin granules (AG) are dictated by the balance between amylopectin synthesis and turnover. Enzymatic activities promoting synthesis include hexokinase (HK), phosphoglucomutases (PGM1/2), UDP-dependent glucose 1 phosphatase (UGPase/AGPase), the commitment step for starch synthesis, Starch/Glycogen synthase (Synthase) and Branching enzyme (BE). Upon synthesis branched amylopectin polymer chains wind expelling water to from the water insoluble starch/ amylopectin granules. Degradation of amylopectin begins with the phosphorylation of the glucose moiety to unwind the amylopectin chain by a glucan kinase allowing access of amylase to the unwound starch molecule releasing glucose. Amylase activity is blocked at sites of glucan phosphorylation necessitating a glucan phosphatase to assure continued degradation. (**C**) PAS and hematoxylin-stained mouse brain histology showing encysted bradyzoites with variability in the distribution of amylopectin. Amylopectin within bradyzoites is evident with the pink stain. Variability of AG within the same cyst and across cysts is evident. Bradyzoite clusters with low levels of amylopectin within the cyst are encircled with a dashed yellow line. (*Scale bar, 10µm*).

Histologically, stored glucans like glycogen (water soluble) and amylopectin, can be detected using Periodic Schiff reagent (PAS), where deposits are evident as a pink stain. In observing PAS-stained tissue cysts within infected mouse brains, we confirmed considerable staining variability in tissue cysts from the same animal (**Fig 1C**). The variations in labeling within tissue cysts confirm that while all the housed bradyzoites are genetically clonal, they follow independent trajectories with regard to AG accumulation. As a histological stain viewed with conventional light transmission microscopy, PAS staining indicates the presence or absence of labeling but lacks, the sensitivity to measure relative levels of AG based on staining. Chemically, the Schiff reagent is rosaniline hydrochloride, which early spectroscopic studies established as brilliantly fluorescent across a broad range of the red spectrum (16). We exploited this property to establish conditions for labeling purified tissue cysts, affording us an optimal signal-to-noise ratio and linearity to capture differences in AG based on PAS staining (see materials and methods).

### Relationship between AG levels and tissue cyst size

Tissue cysts *in vivo* exhibit a broad range of sizes regardless of their time of harvest (5). Historically, tissue cyst size has been used as a measure to assess the effect of mutations, including those associated with AG (21, 22, 28, 29) and pharmacological interventions (30–33). Notably, the loss of the glucan phosphatase TgLaforin does not manifest as a defect in the distribution of tissue cyst size (27). The relationship of AG levels to cyst size was established by measuring the relative intensity following PAS staining of tissue cysts purified at multiple time points (weekly between week 3 and 8 post infection). In pooling multiple timepoints we sought to normalize the effect of the duration of the infection in our assessment. Purified tissue cysts, labeled with FITC conjugated Dolichos lectin (DBA) and PAS were imaged at random with 30 cysts captured at each of the 6 weekly time points post infection. The images were acquired as detailed in the materials and methods with the central slice of a z-stack (0.24μm thickness per slice) in each channel used to determine the mean pixel intensity of the PAS signal and diameter based on DBA demarcating the margins of the cyst using NIH Image J (34). The raw PAS signal without deconvolution was used to determine the mean intensity because deconvolution resulted in the uneven treatment of weak and strong signals to minimize background while accentuating positive signals (data not shown).

As expected, tissue cyst dimensions varied dramatically within the acquired population with sizes ranging from 15-85 μm in diameter (**Fig 2**). Given the overall distribution of sizes we categorized the tissue cysts as small (<30μm), medium (30-60 μm) and large (>60μm) groupings (**Fig. 2**). Notably PAS labeling in each group showed a similar distribution suggesting that tissue cyst size is not an effective predictor of amylopectin levels (**Fig. 2AB**). We therefore examined whether the variability in AG intensity across cysts was driven by the duration of the chronic infection. To address this we developed AmyloQuant, an imaging based application to both quantify and map the relative distribution of AG within tissue cysts based on the intensity of PAS staining.

**Figure 2.**
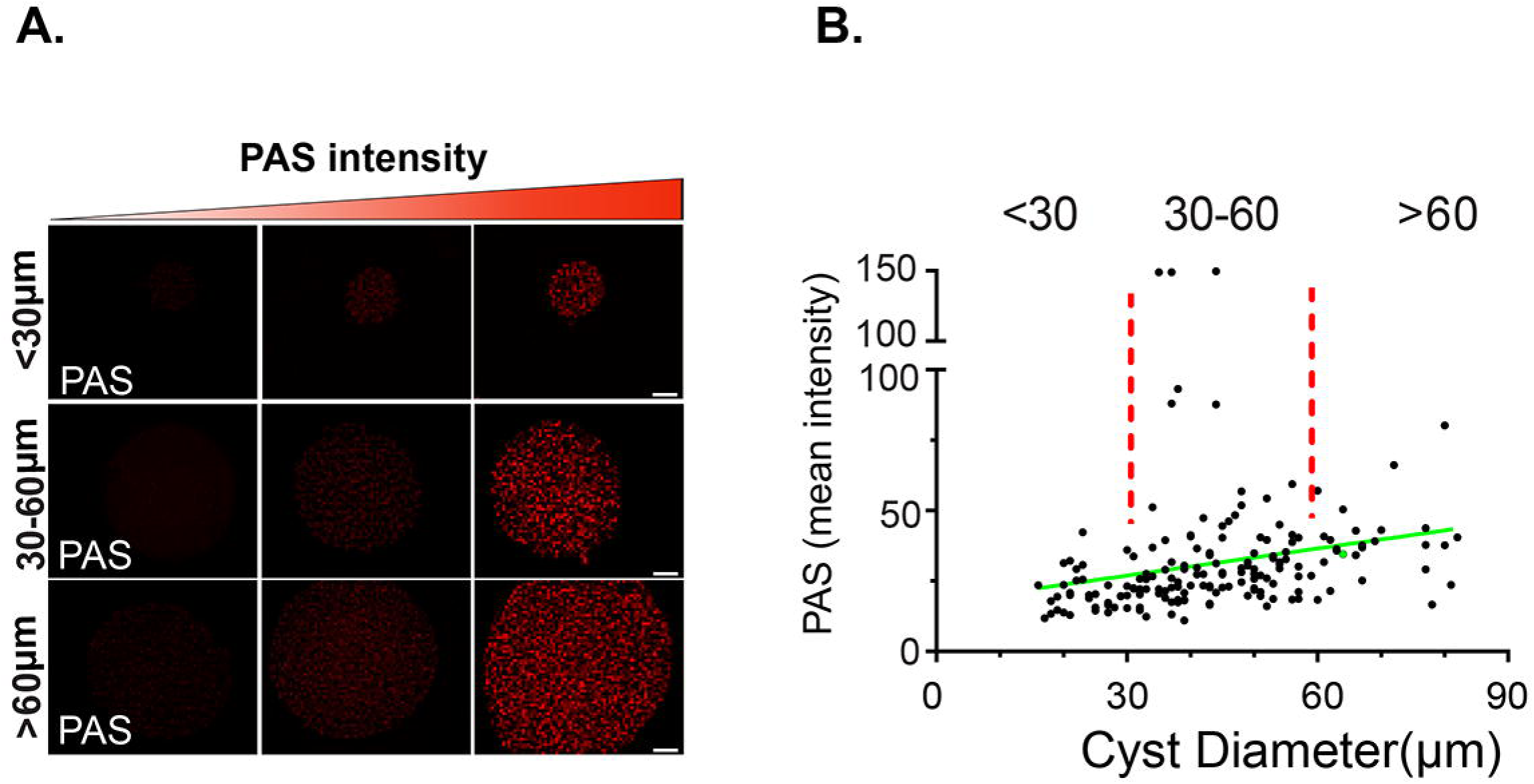
Amylopectin levels are not defined by tissue cyst size. (**A**) Tissue cysts were designated as small (<30μm diameter), medium sized (30-60μm diameter or large (>60μM diameter). Representative images of PAS intensity, defining low, mean and high ranges in cysts of each size class. PAS intensity is in grayscale values determined from the captured grayscale image. Scale bar = 10μm. (**B**) The relationship between cyst size and mean PAS intensity for 180 tissue cysts (30×6) acquired at random at 6 weekly time points from week 3-8 post infection. Of the 180 tissue cysts 38 (21%) were small, 118 (66%) were medium sized and 24 (13%) were large. Tissue cysts in this data set were analyzed by week in Figures 4,5. While a trend line (r2 = 0.0562) suggest a weak correlation between cyst size and AG levels based on PAS intensity. Pearson Coefficient = −0.236.

### Development and implementation of AmyloQuant

The imaging-based application AmyloQuant was developed to establish the intensity of PAS staining and map its distribution within purified tissue cysts. Initial examination of histograms representing the distribution of PAS intensity (AG concentration) revealed that the pattern could be efficiently defined by 4 intensity peaks representing: 1. Background (no labeling), 2. Low intensity pixels, 3. Intermediate intensity pixels and 4. High intensity pixels. The setting of the initial thresholds defining each bin was established using a modified Otsu threshold algorithm (35) to define the specific ranges. The final output from this processing defines the fraction of pixels (relative to total pixels in the imaged volume) belonging to each group. A detailed description of the development of AmyloQuant is presented in the materials and methods.

A graphical user interface (GUI) was developed to facilitate the processing of PAS labeled cyst images (**Fig. 3A**). An outline of the execution of the workflow to capture the levels of AG and their distribution within purified ex vivo cysts is shown in **Fig 3B**. Following the acquisition of images using identical exposure conditions within the experiment (detailed in materials and methods), the AmyloQuant application identifies the location of the cyst within the image defining the region of interest (ROI), which can be manually made larger or smaller using a slider allowing the user to make adjustments to the automatically defined ROI. This isolates the imaged tissue cyst from the surrounding pixels within the captured image, restricting the analysis to the defined ROI. Specific threshold values defining the background, low and moderate thresholds define 4 bin’s allowing for the enumeration of pixels within each bin, from which the total pixel count and proportional pixel count in each class can be determined (**Fig. 3AB**). The effect of altering these thresholds on individual tissue cysts is presented in Supplemental Figure S1, highlighting the importance of standardizing the thresholds to allow for comparative studies.

**Figure 3.**
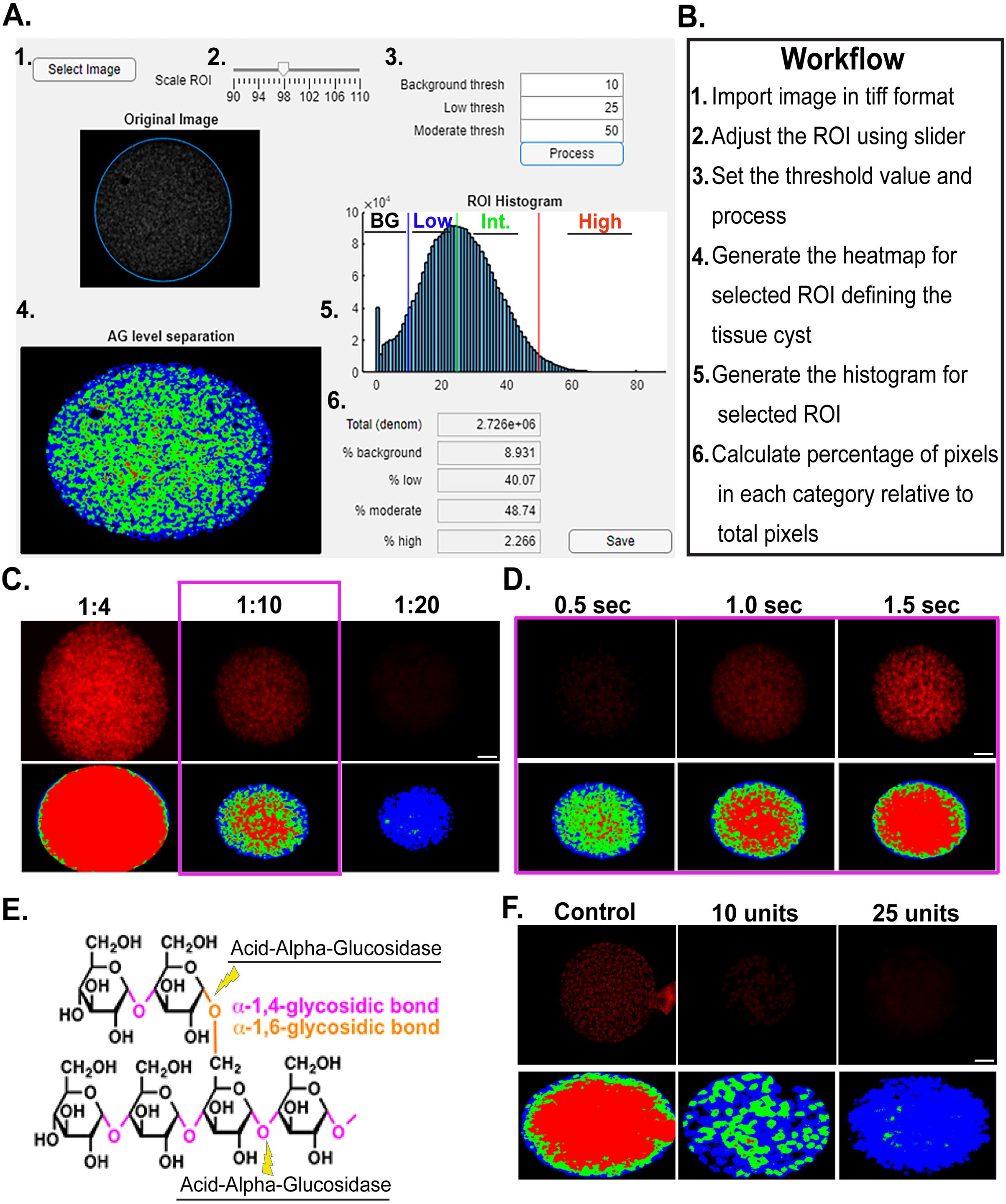
Implementation and validation of AmyloQuant and optimization of PAS labeling. AmyloQuant is an intensity-based image analysis tool that analyzes the heterogeneity in amylopectin distribution in tissue cysts. (**A,B**) The AmyloQuant application allows for the establishment of a region of interest (tissue cyst) within which the distribution of PAS intensity is plotted. The application allows for the setting of 3 threshold cutoffs defining 4 intensity bins, 1. Background (BG-black), Low intensity (Blue), Moderate/Intermediate (Int) intensity (Green) with values above the moderate threshold defining the high intensity pixels (red). The total recorded pixels and proportions in each of the 4 bins is established and a spatial color-coded heat map generated for each cyst. (**C**) Optimization of PAS staining of tissue cyst for analysis. The Schiff reagents in PAS staining were diluted with tap water in a 1:4 (standard protocol), 1:10, and 1:20 ratio. All the images are captured at the same exposure conditions. The top panel shows the cyst image and their respective heatmap from AmyloQuant analysis. Based on the PAS intensity of the image, a 1:10 ratio is selected as the optimum condition. (**D**) Validation of AmyloQuant sensitivity. A PAS-stained cyst was imaged at varying exposure times and analyzed using AmyloQuant with identical threshold values for the pixel bins (BG thresh-10/ Low thresh-25/ High thresh-50). Longer exposure times result in brighter which is reflected in the AmyloQuant generated spatial heat maps. The 1 second exposure was selected as optimal with the optimized PAS staining conditions. (**E**) Acid-Alpha-Glucosidase targets both the linear α1,4 and branched α1,6 glucan linkages resulting in the breakdown of amylopectin. (**F**) The specificity of PAS staining for amylopectin is validated by enzymatic digestion of issue cysts with 10 and 25 units of Acid-Alpha-Glucosidase as is evident from both the intensity following PAS labeling and is captured by the AmyloQuant generated spatial heat map. The control sample was incubated only with a buffer without the addition of acid alpha glucosidase. (*scale bars*,10µm).

Additional standardization was needed by modifying the standard PAS staining protocol, which was developed for largely histological but not fluorescence applications. Our analysis revealed that diluting the Schiff reagent 1:10 provided optimal staining, affording the best dynamic range (**Fig 3C**). Tissue cysts stained under optimal conditions were imaged using different exposure settings. In the example presented we confirm the importance of standardization of exposure conditions as imaging the same cyst with different exposure conditions results in vastly different spatial profiles (**Fig 3D**). Finally, given that PAS can stain glucans other than AG, we employed GAA (Acid-Alpha Glucosidase) treatment to selectively degrade AG and other glucose-based polymers (like glycogen) (**Fig. 3E**). Efficient degradation of AG was noted by the observed loss of PAS labeling that was captured using AmyloQuant (**Fig 3F**).

### The early course of the chronic infection is defined by a slow accumulation of AG

Tissue cysts within the infected animal are highly heterogeneous with regard to the tissue cyst burden, size (5, 36), and replicative state (5). These characteristics vary further the longer the chronic infection persists, which is contrary to the long-prevailing notion that tissue cysts are dormant entities. PAS staining of histological sections from chronically infected brains exhibits qualitative differences (**Fig. 1C**), confirmed by a broad range of mean PAS intensity in purified cysts (**Fig. 2**), suggesting AG levels are dynamic and likely to vary over the course of the infection. Quantification of either absolute or relative AG levels within tissue cysts has not been undertaken. In addition, studies related to AG in the chronic infection have focused on a single time point (typically 4 weeks post infection), with observations inferred to apply the chronic infection in its entirely. To address these issues directly, we leveraged our ability to quantify relative AG levels within tissue cysts using AmyloQuant, tracking them weekly from week 3 to week 8 post-infection, defining the early chronic infection to a more established state.

Tissue cysts purified from infected mouse brains at the indicated time points post infection were deposited on slides, fixed with paraformaldehyde (37), and processed for PAS staining using the optimized protocol. Images were acquired as z-stacks (0.24 microns), with the central stack selected for quantification. Importantly, PAS quantification was performed on the raw image without deconvolution as the application of image deconvolution algorithms (iterative and nearest neighbor) introduced visible artifacts by artificially exaggerating differences in pixel intensity. Cyst images were acquired randomly as they appeared on the slide and processed using AmyloQuant. Analysis, quantifying the number and proportion of pixels in the background range (black-0-10), low (blue-10-25), intermediate (green-25-50), and high ranges (red >50) for each tissue cyst were plotted as a stacked plot (**Fig. 4**). Data from 30 randomly acquired tissue cysts were arrayed in an ordered progression from lowest intensity to highest based on the levels of high-intensity (red) pixels. In order to increase the sensitivity in the high-intensity range, a new set of bins was established in AmyloQuant with the ‘background” level set at a grayscale threshold value of <50 (gray) and the high (red >50) range broken down into grayscale values of 50-75 (light pink), 75-100 (dark pink) and >100 (100-254-purple).

**Figure 4.**
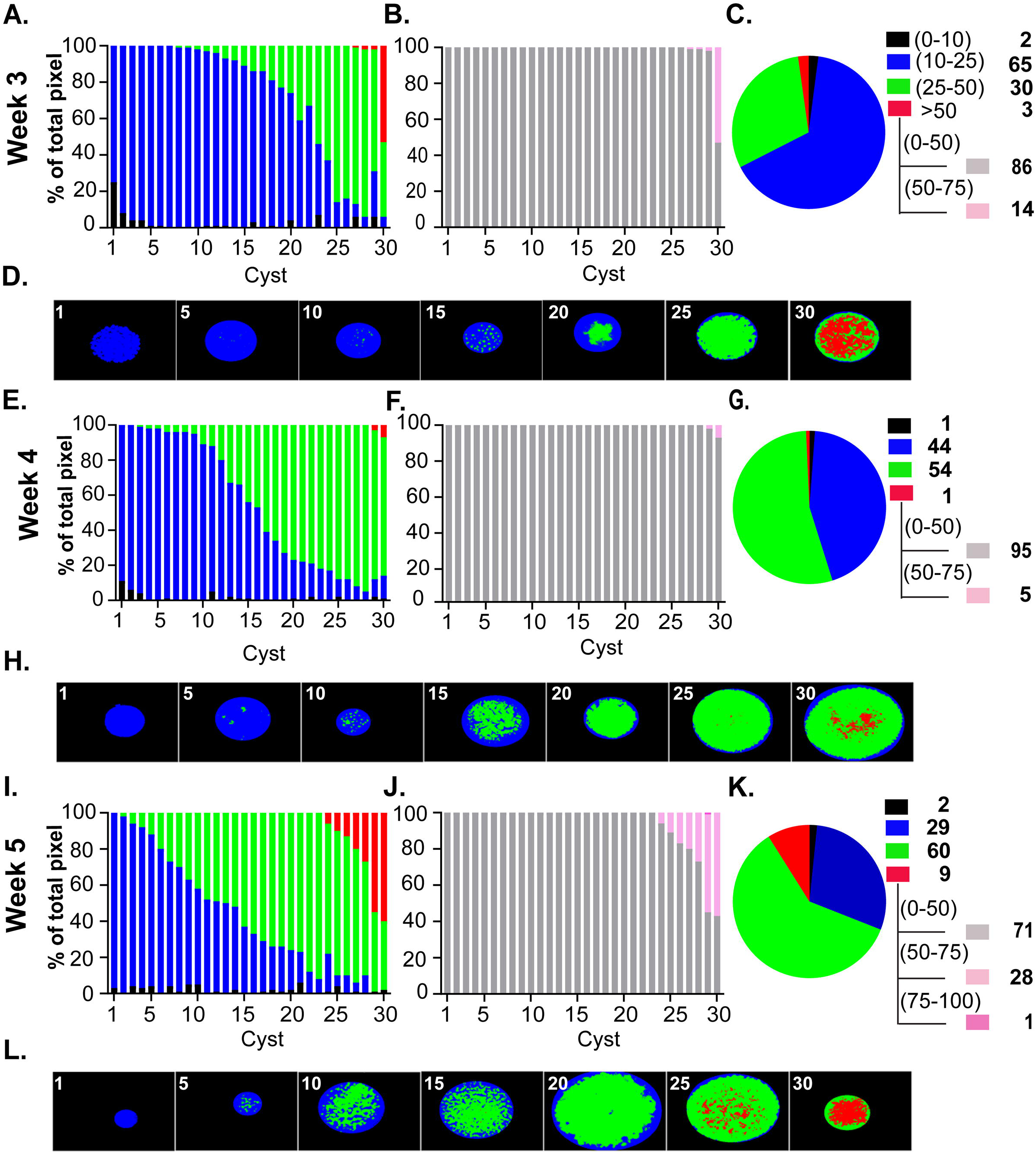
Amylopectin dynamics in the early phase of the chronic infection. PAS stained images of thirty randomly acquired tissue cysts from mice infected for 3 (A,B,C,D), 4 (E,F,G,H) and 5 (I,J,K,L) weeks were analyzed using AmyloQuant. The background (black: 0-10), low (blue:10-25), moderate (green: 25-50) threshold values, with the final high (red: >50) bin defining the 4 bins for analysis presented in a stacked plot format. The tissue cysts were ordered based on the level of high (red) intensity pixels, and secondarily in cysts lacking high intensity pixels on level of moderate (green) intensity pixels. In order to capture the diversity of intensity in high range (red: >50), the background thresholds was set at 50 (gray: 0-50) with high range bins expanded to 50-75 (pink), 75-100 (magenta) and >100 (purple) (B,F,J). The pie charts (C,G,K) represent the pixel intensity distribution following aggregation of the tissue cysts analyzed at each time point. The values, represented as a percentage of the total pixels in each of bin, are presented in the accompanying legend. The tissue cysts represented at each time account for 4258 bradyzoites at week 3, 5485 bradyzoites at week 4 and 5946 bradyzoites at week 5 respectively. Finally, AmyloQuant generated spatial heat maps are presented as thumbnails for every 5^th^ tissue cyst for each week post infection (D,H,L). Consistent with the accumulation of amylopectin during the course of the chronic infection, we observe as general trend of increased AG with a general increase in the levels of intermediate and high intensity pixels reflected for the population in the pie charts (C, G, K) and spatial heat maps (D, H, L).

While tissue cysts can be detected in the brain as early as 2 weeks post-infection, they tend to be underdeveloped, lacking well-defined cyst walls (5). By 3 weeks post-infection, however, tissue cysts can be readily recovered, making this time point widely viewed as the *de facto* onset of the chronic infection (5). Consistent with being populated with bradyzoites, week 3 tissue cysts contain detectable AG with variation in the relative level within tissue cysts accounting for the observed heterogeneity in the population (**Fig. 4A**). In line with recent stage conversion, accumulation of AG within encysted bradyzoites is limited as over two-thirds of the 30 arrayed cysts display low to low-intermediate intensity values for PAS stained pixels (**Fig. 4A**). While relatively uniform with regard to AG levels, 1 cyst out of the 30 displays roughly 50% high-intensity pixels, with others display exceedingly small populations of high-intensity pixels (**Fig. 4A**). Of note, all the high-intensity pixels are within the narrow 50-75 grayscale intensity value range (**Fig. 4B**). To get a sense of AG levels across the population of analyzed week 3 cysts, we combined the values from all 30 cysts and classified them based on the pre-set intensity intervals (**Fig. 4C**). The pie chart for the population confirms the proportions of pixels in each class (**Fig. 4C**) and highlights that most of the high intensity pixels are all in a single tissue cyst with all these pixels confined to the lowest positive interval within the high pixel range (**Fig. 4B**). AmyloQuant generated heat maps showing the distribution of AG based on intensity class within the arrayed sample confirms that intermediate intensity pixels are rare in the 10th cyst and increase in proportion at cyst 15^th^ further increasing to the point where there is a substantial proportion high-intensity pixel (cyst 30/30) (**Fig. 4D**).

As the chronic infection progresses to week 4, we observe a marked shift in the proportion of the intermediate PAS intensity population (green) coming at the expense of low-intensity (blue) pixels. As such, this is consistent with a slow increase in AG accumulation that is likely dictated by the balance between AG synthesis and turnover (**Fig 4E-H**). This population-wide shift is evident, with the intermediate-intensity pixels representing 54% of the population. It is achieved by reducing the proportion of low-intensity pixels relative to week 3. Thus, while there are fewer high-intensity pixels (red), the increase in intermediate-intensity pixels is consistent with either the stabilization of AG levels or, at most, a small increase. Thus, comparing week 3 and week 4, the time points at which most studies on chronic infection are conducted, one might conclude that AG levels within encysted bradyzoites have established an equilibrium.

Interrogation of week 5 derived tissue cysts challenges this assumption as we observe a marked increase in both the number of intermediate level (green) pixels as well as the high intensity (red) pixels. Indeed, 7 of 30 cysts have a significant accumulation of high-intensity pixels such that 9% of the total pixels represent the high-intensity population (**Fig 4I-L**). Overall, the pattern is consistent with a slow but steady increase in AG levels, representing stored metabolic potential potentially reserved for downstream events.

### An unexpected burst in AG accumulation suggests a fundamental metabolic shift

Although our earlier work revealed an increase in the proportion of recently replicated bradyzoites between weeks 5 and 8 (5), a potential role for AG and its stored glucose has not been explored. We, therefore, applied the same methodology to examine tissue cysts harvested at week 6 post-infection (**Fig 5A-D**). Within this population, we observed a marked increase in the proportion of tissue cysts with significant levels of high-intensity pixels (cyst 14/30). This population included tissue cysts that housed >70% of high-intensity pixels, including pixels in the 75-100 and even >100 intensity ranges. On the opposite side of this spectrum, cyst 5/30 presented with only low-intensity pixels, a number that appeared to reverse the trend for accumulation of AG, pointing to the dynamics not being unidirectional. This raises the question of whether processes other than cyst expansion, such as reactivation events, re-seeding events, and resetting the AG level, may occur at this time. Examining the distribution of PAS intensities based on pixel number, a clear shift in the overall population is evident, with both low-intensity and high-intensity populations increasing at the expense of the intermediate population relative to week 5 (**Fig. 4C**). This points to AG dynamics not being unidirectional but rather driven by potentially competing processes. As these studies capture the steady state at each time point for the population, they may reflect distinct physiological demands imposing on AG synthesis and turnover, impacting net accumulation.

**Figure 5.**
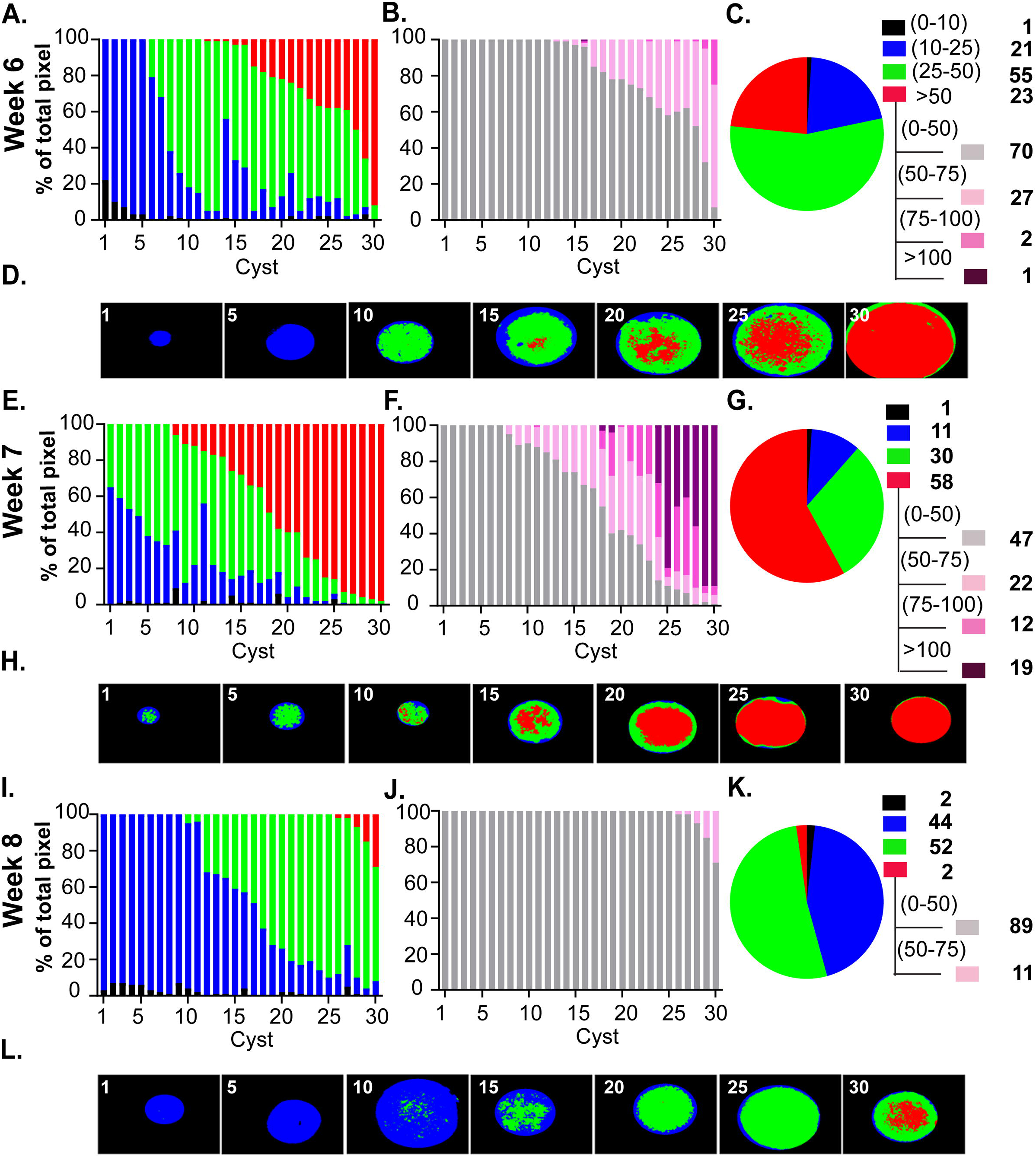
Amylopectin dynamics with the maturation of the chronic infection. PAS stained images of thirty randomly acquired tissue cysts from mice infected for 6 (A,B,C,D), 7 (E,F,G,H) and 8 (I,J,K,L) weeks analyzed using AmyloQuant. The background (black: 0-10), low (blue:10-25), moderate (green: 25-50) threshold values, with the final high (red: >50) bin defining the 4 bins for analysis presented in a stacked plot format. The tissue cysts were ordered based on the level of high (red) intensity pixels, and secondarily in cysts lacking high intensity pixels on level of moderate (green) intensity pixels. In order to capture the diversity of intensity in high range (red: >50), the background thresholds was set at 50 (gray: 0-50) with high range bins expanded to 50-75 (pink), 75-100 (magenta) and >100 (purple) (B,F,J). The pie charts (C,G,K) represent the pixel intensity distribution following aggregation of the tissue cysts analyzed at each time point. The values, represented as a percentage of the total pixels in each of bin, are presented in the accompanying legend. The tissue cysts represented at each time account for 8407 bradyzoites at week 6, 4347 bradyzoites at week 7 and 6802 bradyzoites at week 8 respectively. AmyloQuant generated spatial heat maps are presented as thumbnails for every 5^th^ tissue cyst for each week post infection (D,H,L). The proportion of tissue cysts with high levels of AG increases dramatically at weeks 6 and 7 (A,B, E, F). This is reflected in the distribution of pixel intensities for the bradyzoite population evident in the pie charts (C,G) as well as within the spatial heat maps generated using AmyloQuant (D,E). The transition from week 7 to week 8 is defined by a profound reduction in AG levels (E,F,I,J) withing individual tissue cysts that is reflected in both the overall distribution of PAS intensities evident in the pie charts (G,K) as well as the spatial heat maps bradyzoite population (H,L).

The pattern at week 7 exhibits a profound increase in the accumulation of AG across all cysts examined. Within the pool examined, cyst 21/30 had at least 10 percent of recorded pixels in the high-intensity range (**Fig. 5E-H**). Among these, 5/30 had greater than 90% of pixels in the high-intensity range, which included the majority of pixels in the >100 range. Viewed as a total population, over 58% of pixels were in the high-intensity range, a marked increase from the 23% observed a week prior (**Fig. 5G**). The dramatic accumulation of AG in such a short time window compared to the kinetics across other time points suggests specific priming of a potentially energy-intensive event.

### Rapid utilization of AG and the potential revelation of a programmed cycle

The accumulation of AG in weeks 6 and 7 failed to be sustained. Rather, tissue cysts harvested at week 8 post-infection were marked by a plummeting of overall accumulated AG such that the patterns and distribution across tissue cysts at this time mirrored cysts harvested at week 4 (**Fig. 4E-H**). Taken together, the data suggest the presence of an underlying cycle related to AG that is likely connected to other physiological events, including expansion of tissue cysts, potential disruption and re-seeding of tissue cysts, and elevated levels of potentially synchronized intra-cyst replicative activity.

The data presented in Figures 4 and 5 are from a representative cohort of tissue cysts. In additional studies including tissue cysts fixed with methanol (**Supplemental Fig. S2**) and cysts co-stained with TgIMC3 antibody (**Supplemental Fig. S3**), with 30 cysts labeled at each time point the temporal pattern for TgIMC3 is maintained. While the overall pattern is maintained, we found that fixation followed by storage in methanol (−20°C) replicated the overall pattern with an overall muting of the PAS signal (**Supplemental Fig. S2**). Thus, the temporal pattern for PAS labeling presented in the representative experiments (**Fig. 4,5**) is confirmed with the staining of 90 cysts per time point.

### Correlation of AG dynamics to mitochondrial activity within tissue cysts

As a storage homopolymer of glucose, amylopectin can be viewed as a reserve for energy and biosynthetic needs. The single parasite mitochondrion (38), like all typical mitochondria, is intimately integrated in intermediary metabolism. Fixable mitochondrion targeting reagents like MitoTracker red allow for the selective labeling of active mitochondria within diverse cells, including *Toxoplasma* (39). We established a protocol for the ex-vivo labeling of active mitochondria within freshly isolated tissue cysts (materials and methods). Tissue cysts labeled with MitoTracker display variable levels of labeling (40, 41) that can range from the absence of labeling to very extensive labeling (**Fig. 6A**). Most cysts, however, display a patchwork of labeled mitochondria that do not reveal any specific pattern (**Fig. 6A**). This diversity is notable as it indicates considerable heterogeneity in levels of active mitochondria within tissue cysts and across cysts in the animal. We developed MitoMorph, an imaging-based application to capture and classify mitochondrial forms based on their morphology to capture this diversity (40, 41). In light of not all mitochondria within encysted bradyzoites being active, the application captures and quantified nuclei within the imaged volume, allowing for the number of active (MitoTracker positive) objects relative to the bradyzoite number (number of nuclei) to be established (40, 41).

**Figure 6.**
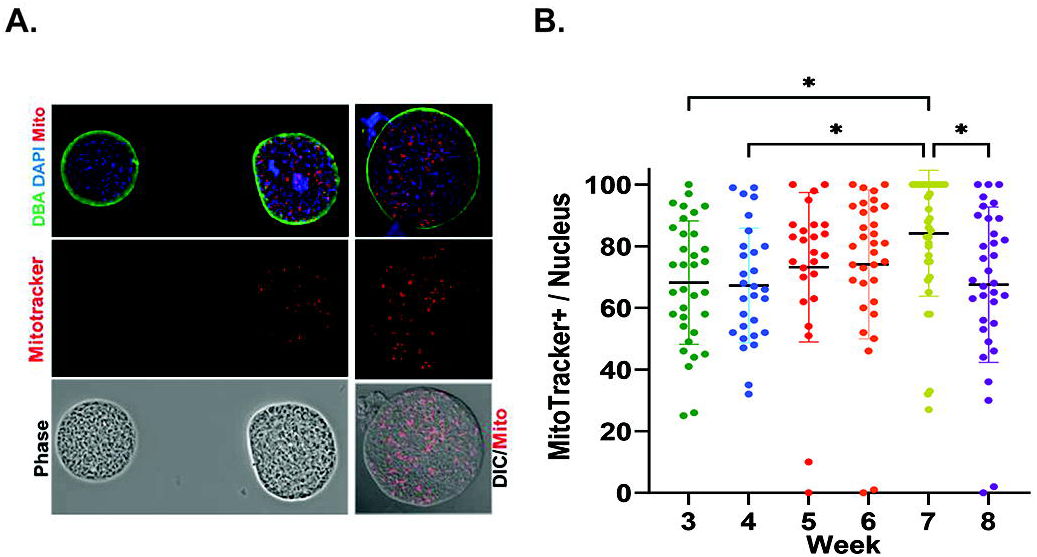
Levels and distribution of active mitochondria based on the membrane potential is highly variable across tissue cysts and displays a temporal profile. **(A)** MitoTracker labeling (red) of freshly isolated ex vivo tissue cysts (DBA-green) reveal considerable heterogeneity ranging from the absence of active mitochondrial to cysts with a high level of activity (left panel). Most tissue cysts display a patchwork of activity as observed in the overlay of the MitoTracker (red) and DIC image (right panel). Nuclei are labeled with DAPI (blue). (**B)** The proportion of active mitochondria relative to nuclei in tissue cysts harvested weekly from week 3 to 8 post infection show a trend of increasing activity from weeks 3-6 followed by a large increase in the number of cysts with >95% of active mitochondria relative to nuclei at week 7. A broad distribution is re-established at week 8 post infection. In instances where the number of mitochondrial profiles exceeded the number of nuclei within the cyst the value was corrected to 100%. Mitochondrial profiles exceeding the number of nuclei could be due to documented fragmentation of the organelle. 1 cyst out of 33 at week 6, 12 cysts out of 39 at week 7 and 2 cysts out of 34 at week 8 were subject to correction. Statistical analysis performed using one-way ANOVA with Tukey’s multiple comparisons test. A *: 0.021-0.030 p-value.

Periodic Acid Schiff reagent has an extremely broad fluorescence emission spectrum (16) that overlaps the emission spectrum for MitoTracker Red (42) and significantly encroaches on the spectrum for MitoTracker Orange (42). Notably, MitoTracker Green and other green emitting potential sensitive dyes that do not spectrally overlap with PAS are not fixable (42), precluding any co-staining. PAS staining was additionally found to interfere with DNA staining dyes like DAPI and Hoechst in a manner that correlated with the overall PAS staining in the sample (**Supplemental Fig. S4**). The effects were variable, resulting in the patchy loss of signal and significant artifactual labeling, as most commonly diffuse staining that made imaging (using BradyCount) (5) as well as manual counting (using labeled counting in Image J) inaccurate and subjective (data not shown). We, therefore, resorted to establishing the levels of active mitochondria across the temporal progression of the chronic infection using matched tissue cysts that were not stained with PAS.

Matched tissue cysts from the same harvests used for PAS staining were labeled with MitoTracker, deposited on slides and fixed prior to staining with Hoescht dye (DNA). Tissue cysts were imaged randomly using fixed image acquisition parameters as a z-stack in both channels. The stack was subjected to deconvolution, and the center slice was used for analysis using MitoMorph (41). As *Toxoplasma* contains a single mitochondrion (38, 39, 43), the ratio of ‘active” (MitoTracker+) objects to total nuclei within the imaged volume served as a surrogate for the proportion of active mitochondria using the specific labeling conditions, acquisition parameters, and thresholds set in MitoMorph (41).

The early phase of the chronic infection (weeks 3-5), which is defined by the slow accumulation of AG (**Fig. 4**), exhibited considerable diversity in the proportion of active mitochondria (**Fig. 6B**). While statistically not different, the mean levels at week 5 are higher than those observed at weeks 3 and 4 (**Fig. 6B**). The proportion of active mitochondria within encysted bradyzoites at week 6 maintains the levels observed at week 5 (**Fig. 6B**). Given the accumulation of AG between weeks 5 and 6 (**Fig. 4I-L and Fig. 5A-D**), it suggests that AG-derived glucose, if involved in driving mitochondrial activity, is generated at a surplus level, driving storage despite utilization. The transition from weeks 6 to 7 is noted by a dramatic statistically significant increase in the proportion of tissue cysts bearing highly energized mitochondria (**Fig. 6B**). While 16% and 15% respectively of week 5 and 6 cysts have > 95% of mitochondria active within the resident bradyzoite population, 47% of week 7 cysts have the same active proportion (**Fig. 6B**). This burst of mitochondrial activity is accompanied by a similar burst in AG accumulation (**Fig. 4,5**). In the absence of a way to directly correlate AG to the mitochondrial activity within individual tissue cysts or bradyzoites due to the spectral overlap, direct correlations to explain this convergence are impossible. However, supporting an integration of AG with overall mitochondrial activity is the observation that the depletion of AG reserves between week 7 and 8 mirrors the effect observed on the proportion of active mitochondria within week 8 cysts-which now mirror the distribution at week 3, suggesting the resetting a cycle for mitochondrial activity.

### Establishing the relationship between AG dynamics and intra-cyst replicative status

The intermediate complex protein TgIMC3 is highly expressed in developing daughter parasites during endodyogeny (44, 45). Once a parasite is “born,” the TgIMC3 signal is reduced to the point that it is no longer detectable in the absence of the re-initiation of a replicative cycle (5). Thus, using quantitative immunofluorescence, the intensity of TgIMC3 labeling can be used to establish the relative ‘age” of bradyzoite populations within tissue cysts. We previously used this metric by measuring the intensity of tissue cysts early (Week 3), in the middle (Week 5), and at later time points (week 8) post-infection (5).

In an ideal setting, measuring TgIMC3 intensity directly in PAS-stained samples would provide a direct correlation between AG levels and the recency of replication. While we could capture TgIMC3 staining in PAS-stained tissue cysts, we found the patterns erratic, especially at time points with high PAS staining (**Supplemental Fig. S3A,B**). We, therefore, undertook TgIMC3 staining of tissue cysts from the same cohort of purified cysts at each time point in the absence of PAS staining and determined the mean TgIMC3 intensity of individual tissue cysts using Image J (5) (**Fig. 6A, B**). These data demonstrate significant differences in TgIMC3 intensity at weeks 4-7, with staining not being different at weeks 3 and 8 when PAS labeling is at its lowest (**Supplemental Fig. S3B**). We, therefore, resorted to addressing the time dependence of TgIMC3 in non-PAS-stained tissue cysts to correlate with prep and time point-matched PAS-stained samples (**Fig. 2,4,5**).

Consistent with our prior findings (5), we observed a marked reduction in the overall TgIMC3 intensity between weeks 3 and 5 (**Fig. 7A, B**). Images of representative high-intensity, mean-intensity, and low-intensity cysts at each time point expose the diversity of the TgIMC3 signal (**Fig 7A**). Interestingly, tissue cysts at week 4 reveal that the bulk of the reduction in the TgIMC3 signal happens between weeks 3-4 and is sustained at week 5. This is entirely consistent with a progression toward a lower replicative state. In our earlier work, we posited that the increase in TgIMC3 intensity between weeks 5 and 8 could be due to a gradual increase across this time frame or, alternatively, an episodic burst of replicative activity and its subsequent stabilization (5). The data are consistent with the latter possibility as we observe a marked increase in mean TgIMC3 intensity across imaged cysts sustained at week 7, settling at an intermediate level between weeks 3 and 5 at week 8, thus matching our prior data (5).

**Figure 7.**
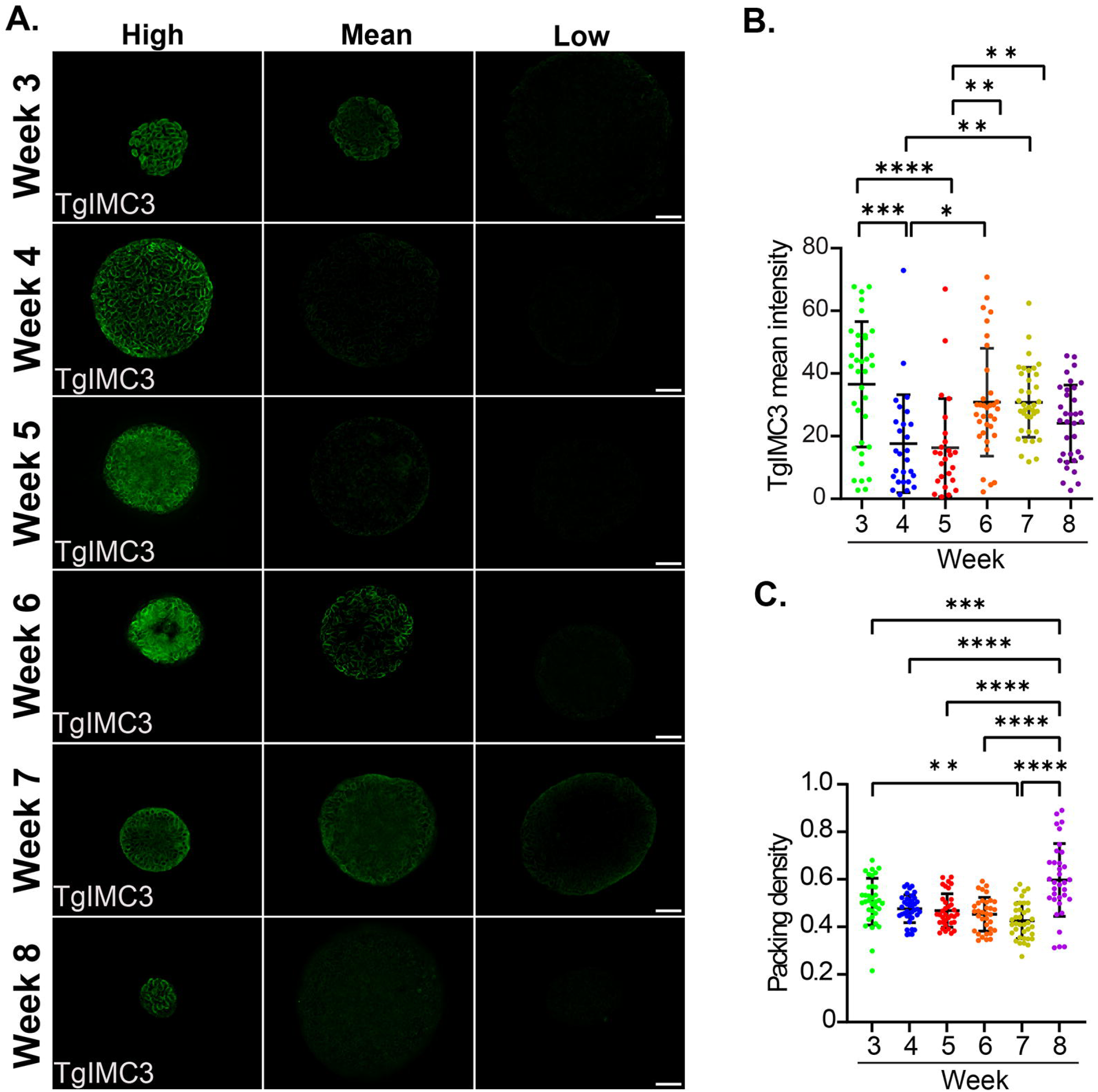
Amylopectin influences the replication of bradyzoites within tissue cysts. (**A**) Recently replicated bradyzoites within tissue cysts can be identified by the intensity of TgIMC3 labeling. The mean intensity of tissue cysts harvested from weeks 3-8 was measured using Image J. The brightest, mean intensity and dimmest cyst at each timepoint Tissue cysts harvested weekly from week 3-8 are represented as a measure of relative mean intensity. All images were captured using identical exposure parameters. Images represent the center slice of a z-stack (z interval of 0.24μm) following iterative deconvolution. Scale bar represents 10 microns. (**B**) Distribution of mean TgIMC3 intensities in tissue cysts harvested at weeks 3-8 reveal an overall pattern of reducing recent replicative activity during the early phase of the chronic infection (week 3-5) that is followed by a burst of recent replication within tissue cysts in the latter half of the cycle (weeks 6-8). Data represent sample means for week 3 (n=35 cysts), week 4 (n=40), week 5 (n=35), week 6 (n=34), week 7 (n=37) and week 8 (n= 37). Statistical Analysis: One way ANOVA with Tukeys multiple comparisons test. Adjusted p values: *: 0.011, **: 0.002, ***: 0.0002, ****: <0.0001 (**C**) The packing density, a measure of cyst occupancy was determined for the TgIMC3 labeled cysts. Packing densities revealed a generally stable pattern from week 3-7 consistent with balanced sporadic intracyst replication. The transition from week 7 to week 8 was noted by extensive intra-cyst bradyzoite replication within most tissue cysts indicative of a coordinated replicative burst. Statistical analysis: One way ANOVA with Tukey’s multiple comparisons test. Adjusted p values: **: 0.0030, ***: 0.0007, ****: < 0.0001.

### Depletion of AG between weeks 7 and 8 correlates with a burst of intra-cyst replication

The packing density, which reflects the occupancy of tissue cysts, provides a measure by which cysts of different sizes can be compared with regard to their relative bradyzoite burden (5). This measure is based on the enumeration of nuclei within the imaged volume (5). In this study, we used a fixed z-step in image acquisition for all cysts regardless of size, allowing for the area encompassed by the margins of the cyst to suffice in determining the packing density.

Due to the interference of PAS with nuclear staining (**Supplemental Fig. S4**), the reliability of determining the packing density of PAS-stained tissue cysts is in question. We, therefore, used the cohort of tissue cysts used for the TgIMC3 intensity (**Fig. 7A,B**) measurements to establish the packing density (PD) distribution from weeks 3-8 (**Fig. 7C**). While the impact of recent replicative events captured with TgIMC3 intensity will, in general, increase the packing density, how it is reflected in the data will depend on the proportion of replicating bradyzoites as well as the actual recency of the event. This effect can, therefore, be muted when measuring the overall mean intensity of TgIMC3 such that low-level replication events may be difficult to resolve whole coordinated recent replication of a substantial proportion of bradyzoites (**Fig. 7AB**).

The packing density distribution while variable showed no statistical difference between weeks 3 and 6 for all pairwise relationships (**Fig. 7C**). The packing density was however statistically significant between week 3 and week 7 purified cysts, but not between the intervening weeks and week 7 (**Fig. 7C**). Cysts at week 7 have the lowest mean PD following a trend from week 3. This pattern is completely reversed between week 7 and 8 resulting in a highly significant increase the PD, indicating a large scale potentially coordinated replication event across tissue cysts within this time interval (**Fig. 7C**). Notable here is that this coordinated burst in intra-cyst replicative activity is temporally connected to the dramatic loss of AG (**Fig. 5**) and overall mitochondrial activity (**Fig. 6**).

## Discussion

Storage glucans like glycogen in animals and starch in plants are used as energy/ metabolic reserves that are charged under conditions of low demand as a resource for future energy and metabolite intensive processes (46–48). In *Toxoplasma*, cytoplasmic amylopectin granules (AG), are more similar to insoluble plant starch than soluble animal glycogen (9). AG are associated with transmission stages, including encysted bradyzoites and sporozoites (4, 8, 9), but not with rapidly replicating tachyzoites, which appear to contain a more labile glycogen-like storage glucan (27). The association of AG with transmission forms had led to the view that their primary, if not sole purpose, is to provide and energy/ metabolite resource to be deployed upon transmission and differentiation into replicative tachyzoites (6, 28). Our work (5), that challenged the long prevailing dogma of bradyzoites lacking replicative potential, presented the possibility that AG could be used during the chronic infection as an energy/ metabolite resource to promote energy intensive processes, including intra-cyst bradyzoite replication.

Studies on the chronic infection (including those focused on the AG pathway) have, and continue to be focused on the tissue cyst, with tissue cyst burden (6, 22, 28, 29, 49) and size (6, 22, 28–30, 49) serving as measures of the extent of the chronic infection. Both measures can be highly variable even in the absence of any genetic or pharmacological intervention making them less than ideal metrics (5, 36). Indeed, following the analysis of a potential relationship between mean AG levels based on PAS intensity and size, we are unable to define any significant correlation functionally linking the two (**Fig. 2**). This points to the need to quantify the distribution of AG within individual cysts, to more robustly and accurately, address AG levels in the progression of the chronic infection.

With our introduction of imaging-based approaches we can effectively deconstruct tissue cysts to the level of encysted bradyzoites, addressing bradyzoite occupancy (5) and mitochondrial activity/morphology (40, 41). In this study we leveraged our experience with imaging-based approaches to address the dynamics of AG during the course of the early and maturing chronic infection following staining with an optimized PAS labeling protocol. We developed AmyloQuant, an imaging-based application allowing for the capture of PAS fluorescence intensity to measure and map relative AG levels within tissue cysts (**Fig. 3**). This application has exposed AG to be considerably more dynamic than previously imagined. First, tissue cysts display considerable heterogeneity in both the levels and distribution of AG at every time point and within every tissue cyst prep (**Fig. 4,5**, **Suppl S2**). This diversity of AG levels within tissue cysts is made more remarkable given that every tissue cyst is genetically clonal having originated from the infection of single parasite. This deviation from uniformity is likely the outcome the loss of replicative synchrony noted in *in vitro* bradyzoites (50, 51) leading to the establishment of distinct physiological trajectories within the same tissue cyst.

Most studies on the chronic infection examine a single time point typically between 3-5 weeks post infection, with analysis restricted to the tissue cyst burden. These studies treat tissue cysts as uniform entities despite clear evidence that they differ greatly in size and bradyzoite occupancy (5). In our earlier work (5), we undertook analyses to determine the progression of the chronic infection which led to the finding that the onset of the chronic infection at week 3 post infection was marked by evidence of considerable replicative activity based on overall labeling with TgIMC3 (5), a marker for the recency of replication (45). The proportion of recently replicated bradyzoites within cysts plummeted by week 5 but showed a partial resurgence at week 8 post infection (5). Building on the skeleton of this potential cycle we determined the relative amylopectin levels within tissue cysts harvested weekly between weeks 3 and 8 using AmyloQuant (**Fig. 3,4,5, Suppl S2**). Not surprisingly, recently formed bradyzoites at the onset of the chronic phase (week 3) contained relatively low levels of AG accumulation (**Fig. 4**). During this phase we observe a steady increase in accumulation of AG with considerable diversity in AG levels between cysts at each time point (**Fig. 4**). This cyst to cyst variability can be partially explained by the fact that stage conversion in the brain is affected by the kinetics of parasite tropism to the brain as well the efficiency of conversion to the encysted state.

In our earlier work, we noted that tissue cysts harvested at week 8 possessed evidence of recent replicative activity that was intermediate that seen at weeks 3 and 5 respectively (5). When we measured the changes in AG levels at weeks 6 and 7 we were surprised by the rapid accumulation of AG initiated at week 6 and progressed through week 7 (**Fig. 5**). Notably the increases in AG levels were not uniform for all cysts, pointing to the physiological and metabolic heterogeneity of the encysted bradyzoites (**Fig. 5**). Despite this heterogeneity however, the overall population of cysts and by extension, the resident bradyzoite population, exposed a potentially pre-programmed response whereby the steady accumulation of AG in weeks 3-5 (**Fig. 4**) is punctuated by a burst of AG storage at weeks 6-7 (**Fig. 5**). Such potentially pre-programmed storage of AG is a likely harbinger for significant energy/metabolic-intensive events. Rapid utilization of AG from high levels at week 7 to levels at week 8 observed early in the chronic infection (weeks 3-4) (**Fig. 4,5**), suggest the resetting of a potential AG-cycle.

The conversion of tachyzoites to bradyzoites in cell culture is associated with a decrease in the mitochondrial membrane potential (52). Imaging of ex vivo tissue cysts labeled with MitoTracker revealed a considerable level of heterogeneity for both mitochondrial activity and morphology (41). This heterogeneity in mitochondrial activity within tissue cysts is evident from the proportion of bradyzoites that label with MitoTracker (**Fig. 6**). Since the parasite contains a single mitochondrion (38), this ratio can be quantified by counting the number of labeled mitochondrial profiles relative to the number of nuclei within the imaged volume (**Fig. 6B**). As expected, the proportion MitoTracker labeled mitochondrial profiles varied significantly from cyst to cyst (**Fig. 6AB**) but showed a generally increasing pattern from week 3-6 (**Fig. 6B**). Coincident with the sharp increase in AG, the transition from week 6 to 7 (**Fig. 5**) correlated with a significant increase in the proportion number of tissue cysts with virtually all mitochondrial profiles active (**Fig. 6**). This suggests a tight functional connection between AG accumulation and mitochondrial activity suggesting with AG accumulation and turnover liberating glucose to drive the membrane potential (ΔΨm). This relationship is reinforced as the apparent utilization AG noted by the precipitous drop in its levels between weeks 7 and 8 (**Fig. 5**) is mirrored with a concomitant decrease in the proportion of active mitochondria within the encysted bradyzoite population (**Fig. 6**).

Such dramatic shifts in both stored energy and energy utilization would only make logical sense if applied to an energy intensive process such as replication. We therefore determined the level of TgIMC3 labeling from week 3-8 (**Fig. 7**). We found that the harshness of PAS treatment had differential effects on TgIMC3 labeling based on AG levels (**Supplemental Fig. S3**). Additional effects were noted with regard to the quality of the DAPI labeling to establish nuclear counts (**Supplemental Fig. S4**). We therefore examined TgIMC3 labeling from week 3-8 on an independent cohort of matched tissue cysts. Consistent with earlier data (5) we observed high TgIMC3 levels at week 3 with a marked decrease at weeks 4 and 5 indicating a marked reduction in recent replication (**Fig. 7B**). The rapid accumulation of AG at weeks 6-7 coincided with an increase in active replication within tissue cysts that was maintained at week 8, at a level between that for weeks 3 and 5 (**Fig. 7B**), thus replicating earlier findings (5).

We next established the packing density of bradyzoites as a function of the progression of the chronic infection in this cohort of tissue cysts. The packing density allows for the comparison of tissue cysts of different sizes (5), with a higher packing density correlating with significant intra-cyst replication. Although there is clear evidence for an increase in recent replication between weeks 5-7 (**Fig. 7AB**), the increases appear to be distributed between tissue cysts in a manner that does not significantly alter the packing density (**Fig. 7C**). This is in sharp contrast to the events between week 7 and 8 which is marked by a more highly synchronized mass replication event resulting in a dramatic increase in the packing density (**Fig. 7C**). This highly coordinated replicative burst of bradyzoites within tissue cysts appears to be the culmination of a build-up of events within which AG plays an important role. Toward comparing the temporal progression of the chronic infection based on the measured physiological parameters, we plotted the mean (+/− SEM) for each parameter as a function of time (week post-infection), separating the early (weeks 3-5) and later (weeks 6-8) time points (**Fig. 8**). Together, the data paint a picture of the dynamics of AG and that the proportion of active mitochondria is very similar, suggesting an integration of AG levels with overall mitochondrial activity (**Fig. 8**). The loss of recent replicative activity between weeks 3-5 correlates with an overall lower level of AG and its slow accumulation (**Fig. 8**). The overall increase in recent replicative activity in the second phase of the cycle is mirrored by the rapid accumulation and subsequent depletion of AG (**Fig. 8**). This replicative phase, while evident with the burst in the packing density at week 8 appears to involve the culmination of events initiated at week 6 as part of a coordinated process (**Fig. 8**).

**Figure 8.**
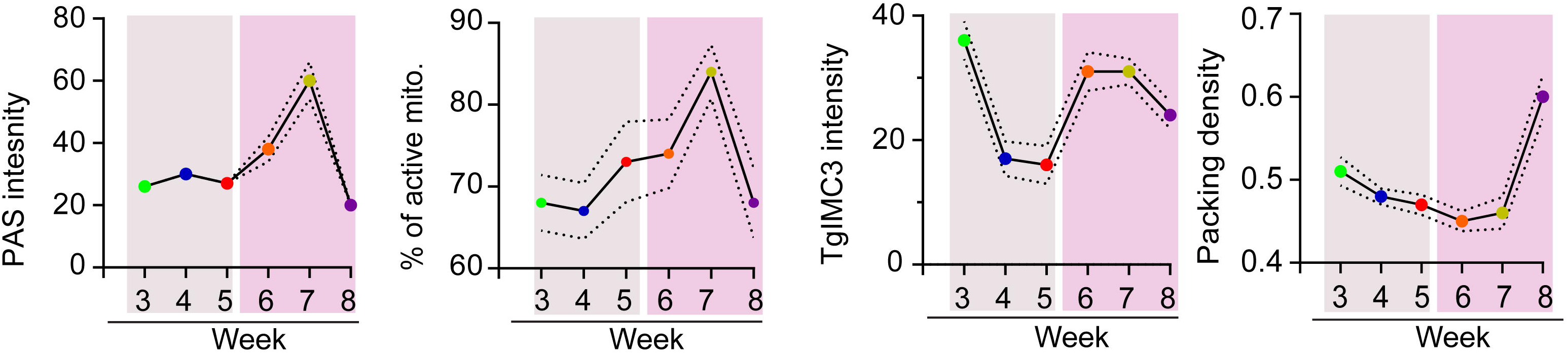
Correlation of AG dynamics, mitochondrial activity, recency of replication and packing density follow distinct patterns. **Panel 1**: Mean PAS intensity for the analyzed tissue cyst populations harvested weekly from week 3-8 post infection reveal a slow increase in the early phase (weeks 3-5) followed by rapid accumulation of AG (weeks 6-7) and subsequent loss at week 8. (Dashed line +/− SEM) **Panel 2**: This pattern is largely mirrored by the proportion of active mitochondria with the increased levels of active mitochondria initiating at week 4, peaking at week 7 post infection. This is followed by a precipitous drop between weeks 7-8 resetting a potential cycle. **Panel 3**: Recency of replication based on mean TgIMC3 intensity reveals a shutting down of active replication between weeks 3-5 a period associated with limited AG accumulation. Increases in AG accumulation at weeks 6-8 are associated with a marked increase (week 6-7) of recent replicative activity that is sustained at week 8. **Panel 4:** A general trend for a stable packing density between weeks 3 and 8 is consistent with balanced cyst expansion matched with sporadic growth. The transition from week 7 to 8 is associated with coordinated intracyst replication that is accompanied by the depletion of AG reserves and overall mitochondrial activity. Statistical analyses: In each case the dashed line represents the boundary of the standard error of the mean (+/− SEM).

In light of the centrality of glucose in intermediary metabolism, a role for stored glucose in the form of AG is not surprising. Its importance is highlighted with the analysis of several mutants involved in both the synthesis and turnover of amylopectin. Interference at the level of AG synthesis, targeting genes commitment step to starch synthesis the UDP-sugar pyrophosphorylase (TgUSP, TgUDP-GPP/ UGPase (TgME49_218200) (21) or Starch/glycogen Synthase (TgSS-TgME49_222800) all result in a lower cyst recovery (22), and further hint toward a defect in reactivation based on cell culture based studies (22). In contrast, deletion of the Starch branching enzyme (TgSBE1-TgME49_316520) failed to have any impact on tissue cyst recovery (53). The cyst burden is dependent on the successful navigation of the acute infection, tropism to the brain, stage conversion and subsequent development in the chronic phase. Thus, any phenotype assessed at the level of the cyst burden can manifest at one or several of these steps. With the typical assessment of phenotypes at 28-30 days post infection (week 4), the true effect of the AG defect may not have yet fully manifested, leaving open the possibility of more significant impacts than currently reported.

Steady state levels of AG are dictated by the effects of both synthesis and degradation. Deletion of starch degrading α-amylase (Tg-αAMY-TgME49_246690) resulted in an overall reduction of the tissue cyst burden (54). The dynamics of AG indicate a level of regulatory control for which the Calcium Dependent Protein Kinase TgCDPK2 (TgME49_225490) has emerged as key component (49). Deletion of this gene results in aberrant accumulation of AG in tachyzoites and within in vitro bradyzoites (49). Loss of TgCDPK2 is reported to eliminate tissue cysts, although these data need to be viewed with caution on account of the low reported cyst yields in animals infected with the wild type Pru strain (49). The protein phosphatase PP2A-holoezyme, has been shown to dephosphorylate TgCDPK2 affecting its activity (55, 56). Ablation of TgPP2A holoenzyme in the ME49 background similarly resulted in the loss of tissue cysts (55, 56). Based on transcriptomic studies, and differential phoshoproteomics, regulation appears to be occurring at the levels of both AG synthesis and turnover (55, 56). This is not surprising for a storage molecule as the overall regulatory environment needs to be programmed for either the “charging” or ‘discharge” of the battery.

Efficient AG degradation is dependent on reversible glucan phosphorylation catalyzed by the kinase-phosphatase pair of the glucan water di-kinase TgGWD (TgME49_214260) (21) and the phosphatase TgLaforin (TgLAF, TgME49_205290) (17, 27). Direct phosphorylation of the glucan chain by GWD facilitates unwinding allowing amylase-mediated degradation to the point of phosphorylation (23, 24, 57) (**Fig. 1B**). TgLaforin, by removing the phosphate from the glucan chain allows for starch degradation to proceed (17, 23). Loss of either TgGWD or TgLaforin is predicted to produce a starch (or glycogen) excess phenotype based on findings from algae to higher plants and animals including mammals (23, 57, 58). Consistent with a role for AG in the chronic infection deletion of TgGWD (21) or TgLaforin (TgLAF) (27) results in a reduction in the overall cyst burden. The effect of the loss of TgLaforin on AG accumulation was found to be nuanced revealing a temporal dimension (27). Notably, the ΔTgLAF mutation did not have a significant impact on AG, based on PAS and AmyloQuant analysis at week 4 post infection (27). In contrast, ΔTgLAF tissue cysts harvested at week 6, displayed a marked increase in the levels of AG accumulation relative to WT or complemented tissue cysts (27). This is notable as at week 6 both an increase in AG levels and recent replication (based on TgIMC3) are evident (**Fig. 8**) pointing to an imbalance caused by the expected reduction in the efficiency of AG turnover. This exaggerated accumulation of AG is reflected in electron microscopic analysis which reveal bradyzoites filled with AG that are enucleated and/or appear dead (27). Furthermore, these data indicate that the AG pathway is potentially druggable, with the effectiveness of potential small molecules being linked to the specific physiological state of bradyzoites along its temporal cycle (17). Not surprisingly, given the emerging importance of the physiological state of encysted bradyzoites is that a clear correlation emerges with regard to the levels of AG and the proportion of ‘active” mitochondria within tissue cysts (**Fig. 4, 5, 6**). Further functional connections of AG to both episodic and synchronized replication events (TgIMC3 intensity) reflected in changes in the packing density point to a central role for AG (**Fig. 7,8**).

A role for stored glucans in the regulation of replicative cycles is evident in the plant kingdom from photosynthetic algae to higher plants (59–62). In general, starch accumulates during the light cycle, but supports anabolic and replicative functions during the dark cycle (59–61). For example, in *Euglena gracilis*, light exposure just before subjective ‘dusk” is most effective in the commitment to cell division, while exposure to light close to subjective ‘dawn” is not (63). Licensing of this commitment is intimately linked to levels of stored starch (63). While light exposure cannot be a factor in dictating the commitment to replication, immuno-metabolic cues associate with the development and maintenance of immune pressure within the host are likely to be sensed and responded to. The transition from tachyzoite to bradyzoites has long been associated with the establishment of the adaptive immune response (reviewed in (64, 65)). More recent work on the maintenance of immune pressure, mediated predominantly by effector T populations and key cytokines like interferon gamma indicates that this response is dynamic (reviewed in (66, 67)). Notable here, is the finding that as the chronic infection progresses, both resident and peripheral T cells exhibit a loss of effector functions correlating with the emergence of suppressive markers of T cell exhaustion (66–69). The emergence of these markers evident from the direct examination of *Toxoplasma* directed T cell populations and transcriptomic analyses (66, 67, 69). This decrease in specific immune pressure coincides with the apparent reprogramming of the AG pathway noted by a rapid accumulation of AG between weeks 5-7 culminating in its apparent utilization between weeks 7-8 (**Fig. 5,8**). With temporal changes in relative mitochondrial (**Fig. 6,8**) and replicative activity (**Fig. 7,8**), suggest an integrated response within encysted bradyzoites.

While the specific triggering signal remains unclear, the apparent temporal changes in immune status are the potential mediators governing AG accumulation and utilization, analogous to light in algae and plants (59–62). As a potential cycle, it is tempting to speculate that spontaneous reactivation events, while controlled, serve to re-invigorate the immune response, potentially tailoring it toward bradyzoite directed antigens thereby establishing a new détente. This new détente could alter the progression through the subsequent temporal cycle, shortening it, lengthening it or maintaining it unchanged. Such an integrated interplay would explain the well established role of the mouse strain on the tissue cyst burden as well as susceptibility to reactivation and associated symptomology during the progression of the chronic infection (70–72). Additionally, it would explain the “life-long persistence” of *Toxoplasma*, with one critical caveat. The encysted bradyzoites that persist are unlikely to be the original parasites establishing the infection but rather their descendants following cycles of reactivation and re-seeding.

In summation, with this study we expose the dynamics of AG accumulation over the first 6 weeks of the chronic *Toxoplasma* infection. In examining this time course, we reveal a previously unrecognized temporal progression that links AG to overall mitochondrial and replicative activities. Unlike tachyzoites bradyzoites demonstrate and high degree of physiological and metabolic heterogeneity that dictates their capacity to replicate (73). These data suggest that AG may serve as a critical metabolic licensing factor, necessitating a minimal level to permit replication of encysted bradyzoites. Thus, while sporadic replication occurs in individual bradyzoites, coordinated large scale replication events require robust coordinated AG accumulation as observed in week 6-7 (**Fig. 5**). The finding that bradyzoites are physiologically heterogenous within and across tissue cysts, yet demonstrate a clear temporal progression exposed by AG dynamics suggests functional entrainment likely co-evolving with the immuno-metabolic status of the host. These findings further reinforce the view that the chronic *Toxoplasma* infection, is an active rather than a dormant phase in the life cycle that simply operates on a longer time scale. This fact needs to be integrated into the design of future studies related to the chronic infection as investigations single time points provide a very limited view, that should not be inferred as applying to the entire life cycle stage.

## Materials and Methods

### Generation and purification of tissue cysts

CBA/J mice of both sexes (Jackson Laboratories, Bar Harbor, ME) were injected intraperitoneally with 20 Type II ME49ΔHXGPRT tissue cysts as previously described (36). Details regarding husbandry of infected animals including monitoring for symptoms and euthanasia were conducted as previously described (5, 36, 37). Tissue cysts were purified at weeks 3,4,5,6,7, and 8 post infection. In each instance purifications were done two days following the completion of the week (for example: Week 3: day 23). All protocols were carried out under the approval of the University of Kentucky’s Institutional Animal Care and Use Committee (IACUC).

### Tissue cyst purification

Tissue cysts were recovered from infected brains at the indicated time points with 2 brains being processed for each prep as described in detail (37). Purified tissue cysts were obtained from the infected brain homogenate using a discontinuous Percoll gradient and quantified following fractionation of as previously described (37). Recovered tissue cysts were pelleted onto glass slides (200-300 cysts/slide) and fixed in either 4% paraformaldehyde in PBS (PFA) for 30 minutes or in absolute methanol at −20°C as indicated. Paraformaldehyde fixed tissue cysts on slides were stored in PBS at 4°C while methanol fixed slides were stored in absolute methanol at −20°C prior to labeling.

### Histology of infected mouse brains

Brains from chronically infected mice were harvested intact and fixed in Neutral Buffered Formalin (10% formalin in PBS) for 24 hours and stored in 70% Ethanol. Individual hemispheres were paraffin embedded and 5μm sections generated and fixed on slides. Sections were sequentially stained with hematoxylin and PAS at the Biospecimen Procurement and Translational Pathology Shared Resource Facility at the University of Kentucky Markey Cancer Center using standard protocols. Stained sections were acquired using a Zeiss Axioscan Z1/7 slide scanner and analyzed using Zeiss Zen software.

### <u>P</u>eriodic <u>A</u>cid-<u>S</u>chiff (PAS) labeling of tissue cysts

The tissue cysts purified from the infected mouse brain were centrifuged and deposited on a glass slide using a Cytospin centrifuge (37). The cyst deposited on the glass slide was fixed either in methanol (100%, stored at −20 °C) or in paraformaldehyde (PFA) (4% in phosphate-buffered saline (PBS). Tissue cysts fixed in PFA were permeabilized in 0.1% triton solution in PBS^++^((PBS containing 0.5 mM MgCl_2_) for 10 minutes. The tissue cysts fixed in methanol were proceeded for PAS staining. The sample was washed thrice in tap water in a caplin jar and incubated with 0.5% of periodic acid (Newcom Supply, cat# 13308A) for exactly 5 minutes. For the optimized labeling protocol Schiff’s reagent (Newcom Supply, Cat# 1371A) was diluted 1:10 in tap water and added to the slide in the dark for exactly 10 minutes. Other dilutions tested were 1:4 (Manufacturers recommendation) and 1:20. The slides were transferred to a Coplin jar and washed 10 times with tap water. Slides were subsequently counterstained with Hoescht dye to image nuclei. As noted in the results, the harsh conditions of Periodic-Schiff regent caused interference with the labeling of some antibodies including a rat anti-TgIMC3 antibody (Donkin et al. in preparation). In addition, high intensity PAS labeling resulted in the distortion of the nuclear staining patterns. For these reasons, the labeling and quantification of these signals was performed in tissue cysts from the same preps that were not PAS labeled. Notably staining with *Dolichos biflorus* lectin was not impacted by PAS.

### MitoTracker labeling of *ex vivo* tissue cysts

Pooled Percoll fractions generated using the standard cyst purification protocol (37) containing a minimum of 2500 tissue cysts were diluted 1:5 with Artificial Intracellular Salts Buffer (AISS) (74) and the cysts pelleted for 5 minutes at 2000 rpm at ambient temperature in a sterile 15ml conical tube. The supernatant was aspirated leaving roughly 250μl into which the tissue cyst pellet (that may not be clearly visible) was resuspended. 2.75ml of Prewarmed AISS buffer with 25nM MitoTracker was added and the tube incubated within a pre-warmed water bath in the incubator for 45 minutes. The tube was mixed every 15 minutes. Following this incubation 12ml of prewarmed AISS was added to chase the MitoTracker and incubated for an additional 30 minutes. The labeled tissue cyst were deposited on glass slides using a Cytospin centrifuge as previously described and fixed in 4%PFA (37), prior to storage as described above. Cyst nuclei were labeled using Hoescht dye prior to imaging as described below.

### Immunofluorescence labeling

PFA fixed tissue cysts deposited on slides were permeabilized with 0.2% Triton X-100 in PBS, washed and labeled with an affinity purified rat anti-TgIMC3 antibody (1:250 dilution in PBS with 3%BSA for 1 hour at room temperature). Following 3 PBS washes slides were labeled with goat anti-rat conjugated with Oregon Green (1:2000, Molecular Probes/ Thermo) and Hoescht dye (300nM, Molecular Probes/ Thermo). Stained tissue cysts were sealed under a coverslip using MOWIOL prior to imaging. Staining of the tissue cyst wall was performed using FITC-conjugated Dolichos biflorus lectin (DBA, Vector Laboratories) at a 1:2000 dilution.

### Amyloglucosidase digestion

Glass slide with deposited purified tissue cysts were washed thrice in 1X citric acid buffer (100mM citric acid/ 100mM dibasic sodium citrate sesquihydrate pH4.6-Sigma) in a Coplin jar. 10 or 25 units of Amyloglucosidase from *Asperigillus niger* (EC-232-844-2)(Megazyme, E-AMGDF, cat# A7096) enzyme were diluted in 1X citric acid buffer and transferred directly onto the site of the deposited tissue cysts onto the glass slide with the region containing tissue cysts outlined using a wax pencil. The control slides were treated with 1X citric acid buffer. Both control and test slides were incubated at room temperature overnight in a tightly sealed, moist chamber. The slides were washed 10 times in 1X PBS++ (PBS with 0.5mM Ca/Mg) buffer in a Coplin jar and processed for PAS and Hoescht dye staining prior to imaging.

### Image acquisition and analysis

The images were acquired on a Zeiss AxioVision upright microscope using a 100X 1.4NA oil immersion objective. Images were acquired as Z-stacks centered around the center of the imaged tissue cyst, using a Zeiss AxioCam MRM digital camera using Zeiss Zen software to control both the camera and motorized stage. The number of Z-stacks acquired was dependent on the size of the thickness of the tissue cyst being imaged although the height of the z-interval was kept constant at 0.24 μm. Acquisition of discrete entities (nuclei, mitochondria and TgIMC3 scaffolds) were subjected to deconvolution using the iterative algorithm in the Zeiss Zen software. For the quantification of PAS staining deconvolution of the signal was not used as the application of the software artificially “normalized” the signal intensity by preferentially exaggerating weak signals thereby reducing the dynamic range. Images in all channels (Nuclei-blue, TgIMC3-FITC-green, and PAS-Texas Red were acquired as co-registered grayscale images using fixed exposure times that were empirically determined to provide the optimal signal to noise profile for each channel independently. As PAS treatment was observed to affect the detection of TgIMC3 (**Suppl S3**), all quantification of TgIMC3 was performed on samples that were not PAS stained. The center slice from the z-stack image of each cyst (either deconvoluted or not) was selected and exported in a TIFF format without compression for further analysis. The cyst images with an even number of slices in the Z stack, the slice below the center, were selected. Quantification of AG levels based on PAS staining using AmyloQuant was performed as below with the workflow outlined in **Figure 3B**. Quantification of TgIMC3 intensity within individual tissue cysts was established using Image J as described previously (5).

### Packing density

Tissue cysts were co-stained with DAPI, and DBA-FITC was used to label the bradyzoites nucleus and highlight the cyst periphery by labeling the cyst wall. The cyst diameter was determined in Fiji (NIH-Image J) (34) by drawing the ROI around the cyst, using DBA staining as the reference. The Multi-point counter mark and count feature in NIH-Image J was used to manually count nuclei within the imaged volume. Once the value of cyst diameter and total nuclear number was obtained, packing density was calculated using the formula describing this calculation is PD = N / (πr^2^ x h), where PD=packing density, N = # of bradyzoites/ nuclei, r = cyst radius, h=height of selected slices which is constant (0.24 µm), and represents the z-stack interval.

### Statistical Analyses

Statistical analyses were performed using GraphPad with specific tests applied in the figure legends.

## Development of AmyloQuant

The images for AG analyses were processed using the following algorithm. From the 8-bit grayscale image AGo, first, a modified Otsu threshold (35) for intensity (to) was obtained, the threshold was scaled by a factor (<1), and the scaled threshold was used to binarize the image. The binary image and “region props” function in Matlab was used to detect the boundary of the cyst. This boundary generated a circular mask delineating the region of interest (ROI). The detected ROI was visualized, and a manual correction factor α (selected via the user interface) was applied, αROI (making the ROI larger or smaller) when necessary. The original grayscale image AG_o_ was multiplied by the mask to set all pixels outside of the ROI to zero, and then the image was segmented into 4 binary images using three pre-defined intensity cutoffs, i_m_, i_l_, and i_b_ as follows: AG_h_ was computed from AG_o_ using a condition that all pixels within AG_o_ with intensity values ≥ i_m_ were set equal to 1 and all pixels with intensities < i_m_ were set to be 0. Likewise, AGm was computed by setting all pixels with intensities ≥ il and <im to be equal to 1 and the rest equal to 0 and for AGl, the pixels with intensities ≥ ib and <il was set equal to 1, and the rest were set to 0. All pixels within the ROI with intensities <i_b_ were set to be equal to 1 and others to be 0 to generate AG_b_. The total number of pixels with values equal to 1 were counted within each of the four images to generate counts k_h_, k_m_, k_l_, and k_b._ These counts were then normalized by k_t_, the total number of pixels within the ROI, e.g., k sub h over k sub t, and expressed as a percent fraction of the image intensity within each of the four intensity ranges.

AG_h_(x,y) = 1 if AG_o_(x,y) ≥ i_m_ else AG_h_(x,y) = 0

AG_m_(x,y) = 1 if i_l_ ≤ AG_o_(x,y) < i_m_ else AG_m_(x,y) = 0

AG_l_(x,y) = 1 if i_b_ ≤ AG_o_(x,y) < i_l_ else AG_l_(x,y) = 0

AG_b_(x,y) = 1 if AG_o_(x,y) < i_b_ else AG_b_(x,y) = 0

The user interface for the program includes a slider labeled ‘Scale ROI,’ which allows for the selection of the α value, allowing for adjustments ranging from 90% to 110% of the automatically detected ROI. The pre-defined intensity cutoffs of i_b_, i_l_, i_m_ are input into the ‘Background thresh,’ ‘Low thresh,’ and ‘Moderate thresh’ fields, respectively (**Fig. 3A**). The pixels present within the ROI can be visualized in the histogram displayed along with vertical lines representing the cutoffs used. The 4 binary images created are merged into a single image showing all low-intensity pixels in blue, moderate-intensity pixels in green, and high-intensity pixels in red, with the background being black. The percent fraction of the intensities within each of the four intensity ranges and the total number of pixels within the ROI are displayed. The code for the application was developed using Matlab.

## Acknowledgements

The authors acknowledge the support and assistance of the staff of the vivarium from the Department of Animal Laboratory Medicine at the University of Kentucky. We thank Dr. Matthieu Colpaert for his insights regarding the evolution of amylopectin metabolism in algae and plants. This work is supported by grants from the US National Institutes of Health RO1AI145335 (to AP and APS) and R21 AI150631 awarded to APS. RDM was supported by GRFP 1247392 from the NSF.

## Author Contributions

Conceptualization (AT, AP and APS), Methodology (AT, JSM, AP and APS), Formal analysis (AT, JSM, RWD, RDM, AP and APS), Investigation (AT, JSM, RWD, RDM, AP and APS), Writing-original draft (AT, APS), Writing-review and editing (AT, JSM, RWD, RDM, AP and APS), Discussions (MSG), Visualization (AT, JSM, AP, APS), Project administration (APS), Funding acquisition (AP, APS).

## Conflicts of Interest

None

## Supplemental Data Figure Legends

**Supplemental Figure S1. Effect of AmyloQuant threshold value settings on the distribution of PAS intensities.** Application of distinct threshold values in AmyloQuant in the same tissue cyst image (original images are adjusted) results in a fundamentally different patterns to define the background, low, intermediate and high bins thus impacting the representation of AG distribution. The optimized setting (Intermediate) of the threshold value provided the most balanced representation for typical tissue cysts. Thumbnails below the intensity histograms present the AmyloQuant generated heat maps for each of the threshold setting presented here.

**Supplemental Figure S2. Amylopectin dynamics revealed in methanol fixed tissue cyst following PAS staining and analysis in AmyloQuant** analysis of tissue cysts fixed in methanol. The bar graphs represent the ordered distribution of intensities in the background, low, intermediate and high ranges following the setting of bins at : BG: 0-10 (black), Low10-25 (blue), Intermediate 25-50 (green) and high >50 (red). The heatmaps under each set of bar graphs represent AmyloQuant generated spatial distributions of the 30 tissue cysts at 5 cyst intervals. While the overall AG pattern from week 3-8 is identical to that observed for PAS labeling of over this temporal course, a significant loss of signal is evident for methanol fixed tissue cysts. The amylopectin distribution pattern in chronic infection’s early and late phases is similar to the PFA fixed and unaffected, irrespective of the fixation condition.

**Supplemental Figure S3. Interference of PAS staining with TgIMC3 labeling. (A)** Counter staining of PAS stained tissue cysts affected the efficiency of TgIMC3 labeling. In general poor TgIMC3 staining was noted in tissue cysts with high PAS labeling. Scale bar represents 10 μm. (**B**) Head to head comparison of TgIMC3 labeling on both untreated (-PAS) and PAS treated (+PAS) tissue cysts shows no difference at time points with low AG (week 3 and 8), but significant PAS dependent differences in TgIMC3 labeling at time points associated with higher levels of AG and thus higher PAS binding. Statistical Analysis: One way ANOVA with Mann-Whitney Test. P values: ns: not significant, **: 0.0047, ****= < 0.0001.

**Supplemental Figure S4. PAS staining impacts the quality of DAPI labeling compromising the ability to accurately count nuclei**. (**A**) PAS labeling of PFA and methanol (not shown) fixed tissue cysts affected the integrity and quality of nuclear staining that resulted in differentially diffuse nuclear profiles. While not uniform, the effect on PAS on nuclear staining was markedly more pronounced in tissue cysts from time points associated with higher levels of AG and thus PAS labeling. Additional factors leading to the exclusion of tissue cysts for analysis included non circular cysts and clearly damaged broken cysts. These issues precluded the accurate demarcation of the whole tissue cyst in AmyloQuant affecting the accuracy of both PAS and nuclear quantification. (**B**) The proportion of PAS/DAPI co-stained tissue cysts within which the number of nuclei could not be accurately counted was significantly greater at time points associated with high PAS labeling (weeks 5-7). For this reason, measurements relating to the packing density which is dependent in the accuracy of the nuclear count was performed on non-PAS staining tissue cyst from the same cohort at each time point. Damage to nuclei is likely connected to the low pH associated with deposited Schiff periodic acid dye in an AG concentration dependent manner.

## Notes

### Competing Interest Statement

The authors have declared no competing interest.

### Summary of Updates

Addition of author Matthew S. Gentry Correction of typographical and grammatic mistakes and clarification of an experimental protocol. No changes in data and outcomes/ interpretation

## Literature Cited

1. J. P. Dubey, J. L. Jones, *Toxoplasma gondii* infection in humans and animals in the United States. International journal for parasitology 38, 1257–1278 (2008).

2. D. E. Hill, S. Chirukandoth, J. P. Dubey, Biology and epidemiology of *Toxoplasma gondii* in man and animals. Anim Health Res Rev 6, 41–61 (2005).

3. A. P. Sinai et al., Reexamining Chronic *Toxoplasma gondii* Infection: Surprising Activity for a “Dormant” Parasite. Curr Clin Microbiol Rep 3, 175–185 (2016).

4. J. P. Dubey, D. S. Lindsay, C. A. Speer, Structures of *Toxoplasma gondii* tachyzoites, bradyzoites, and sporozoites and biology and development of tissue cysts. Clin Microbiol Rev 11, 267–299 (1998).

5. E. Watts et al., Novel Approaches Reveal that *Toxoplasma gondii* Bradyzoites within Tissue Cysts Are Dynamic and Replicating Entities In Vivo. mBio 6, e01155–01115 (2015).

6. V. Tu, R. Yakubu, L. M. Weiss, Observations on bradyzoite biology. Microbes and infection / Institut Pasteur 20, 466–476 (2018).

7. T. V. Beyer, C. Siim, W. M. Hutchison, [Cytochemical study of different stages in the life cycle of *Toxoplasma gondii*. I. Amylopectin and lipids in endozoites]. Tsitologiia 19, 681–685 (1977).

8. A. Coppin et al., Developmentally regulated biosynthesis of carbohydrate and storage polysaccharide during differentiation and tissue cyst formation in *Toxoplasma gondii*. Biochimie 85, 353–361 (2003).

9. A. Coppin et al., Evolution of plant-like crystalline storage polysaccharide in the protozoan parasite *Toxoplasma gondii* argues for a red alga ancestry. J Mol Evol 60, 257–267 (2005).

10. Y. Guerardel et al., Amylopectin biogenesis and characterization in the protozoan parasite *Toxoplasma gondii*, the intracellular development of which is restricted in the HepG2 cell line. Microbes and infection / Institut Pasteur 7, 41–48 (2005).

11. L. Lim, G. I. McFadden, The evolution, metabolism and functions of the apicoplast. Philos Trans R Soc Lond B Biol Sci 365, 749–763 (2010).

12. S. Ball, C. Colleoni, U. Cenci, J. N. Raj, C. Tirtiaux, The evolution of glycogen and starch metabolism in eukaryotes gives molecular clues to understand the establishment of plastid endosymbiosis. J Exp Bot 62, 1775–1801 (2011).

13. L. M. Weiss, K. Kim, The development and biology of bradyzoites of *Toxoplasma gondii*. Front Biosci 5, D391–405 (2000).

14. D. J. Ferguson, Use of molecular and ultrastructural markers to evaluate stage conversion of *Toxoplasma gondii* in both the intermediate and definitive host. International journal for parasitology 34, 347–360 (2004).

15. D. J. Ferguson, W. M. Hutchison, An ultrastructural study of the early development and tissue cyst formation of *Toxoplasma gondii* in the brains of mice. Parasitology research 73, 483–491 (1987).

16. P. J. Stoward, Studies in fluorescence histochemistry. II. The demonstration of periodate-reactive mucosubstances with pseudo-Schiff reagents. J R Microsc Soc 87, 237–246 (1967).

17. R. D. Murphy et al., The Toxoplasma glucan phosphatase TgLaforin utilizes a distinct functional mechanism that can be exploited by therapeutic inhibitors. The Journal of biological chemistry 298, 102089 (2022).

18. P. L. Keeling, A. M. Myers, Biochemistry and genetics of starch synthesis. Annu Rev Food Sci Technol 1, 271–303 (2010).

19. E. V. Quach et al., Phosphoglucomutase 1 contributes to optimal cyst development in *Toxoplasma gondii*. BMC Res Notes 15, 188 (2022).

20. S. Saha, B. I. Coleman, R. Dubey, I. J. Blader, M. J. Gubbels, Two Phosphoglucomutase Paralogs Facilitate Ionophore-Triggered Secretion of the Toxoplasma Micronemes. mSphere 2 (2017).

21. P. Chen et al., Key roles of amylopectin synthesis and degradation enzymes in the establishment and reactivation of chronic toxoplasmosis. Anim Dis 3 (2023).

22. C. Lyu et al., Role of amylopectin synthesis in *Toxoplasma gondii* and its implication in vaccine development against toxoplasmosis. Open Biol 11, 200384 (2021).

23. M. Stitt, S. C. Zeeman, Starch turnover: pathways, regulation and role in growth. Curr Opin Plant Biol 15, 282–292 (2012).

24. J. Compart, A. Apriyanto, J. Fettke, Glucan, water dikinase (GWD) penetrates the starch granule surface and introduces C6 phosphate in the vicinity of branching points. Carbohydr Polym 321, 121321 (2023).

25. M. Hejazi et al., Glucan, water dikinase phosphorylates crystalline maltodextrins and thereby initiates solubilization. Plant J 55, 323–334 (2008).

26. M. S. Gentry et al., The phosphatase laforin crosses evolutionary boundaries and links carbohydrate metabolism to neuronal disease. J Cell Biol 178, 477–488 (2007).

27. R. D. Murphy et al., TgLaforin, a glucan phosphatase, reveals the dynamic role of storage polysaccharides in Toxoplasma gondii tachyzoites and bradyzoites. bioRxiv 10.1101/2023.09.29.560185 (2023).

28. M. Pan et al., The determinants regulating *Toxoplasma gondii* bradyzoite development. Front Microbiol 13, 1027073 (2022).

29. T. Sugi, V. Tu, Y. Ma, T. Tomita, L. M. Weiss, *Toxoplasma gondii* Requires Glycogen Phosphorylase for Balancing Amylopectin Storage and for Efficient Production of Brain Cysts. mBio 8 (2017).

30. A. M. Sullivan et al., Evidence for finely-regulated asynchronous growth of *Toxoplasma gondii* cysts based on data-driven model selection. PLoS computational biology 9, e1003283 (2013).

31. N. Anghel et al., Endochin-like quinolones (ELQs) and bumped kinase inhibitors (BKIs): Synergistic and additive effects of combined treatments against Neospora caninum infection in vitro and in vivo. Int J Parasitol Drugs Drug Resist 17, 92–106 (2021).

32. J. S. Doggett et al., Endochin-like quinolones are highly efficacious against acute and latent experimental toxoplasmosis. Proceedings of the National Academy of Sciences of the United States of America 109, 15936–15941 (2012).

33. R. S. Vidadala et al., Development of an Orally Available and Central Nervous System (CNS) Penetrant *Toxoplasma gondii* Calcium-Dependent Protein Kinase 1 (TgCDPK1) Inhibitor with Minimal Human Ether-a-go-go-Related Gene (hERG) Activity for the Treatment of Toxoplasmosis. Journal of medicinal chemistry 59, 6531–6546 (2016).

34. J. Schindelin et al., Fiji: an open-source platform for biological-image analysis. Nature methods 9, 676–682 (2012).

35. N. Otsu, A threshold selection method from gray-level histograms IEEE Transactions on Systems Man and Cybernetics 9, 62–66 (1979).

36. C. A. Troublefield, J. S. Miracle, R. D. Murphy, R. W. Donkin, A. P. Sinai, Factors Influencing Tissue Cyst Yield in a Murine Model of Chronic Toxoplasmosis. Infection and immunity 91, e0056622 (2023).

37. E. A. Watts, A. Dhara, A. P. Sinai, Purification *Toxoplasma gondii* Tissue Cysts Using Percoll Gradients. Curr Protoc Microbiol 45, 20C 22 21-20C 22 19 (2017).

38. F. Seeber, D. J. Ferguson, U. Gross, *Toxoplasma gondii*: a paraformaldehyde-insensitive diaphorase activity acts as a specific histochemical marker for the single mitochondrion. Experimental parasitology 89, 137–139 (1998).

39. J. Ovciarikova, L. Lemgruber, K. L. Stilger, W. J. Sullivan, L. Sheiner, Mitochondrial behaviour throughout the lytic cycle of *Toxoplasma gondii*. Scientific reports 7, 42746 (2017).

40. B. C. Place, C. Troublefield, R. D. Murphy, A. P. Sinai, A. Patwardhan, Computer Aided Image Processing to Facilitate Determination of Congruence in Manual Classification of Mitochondrial Morphologies in *Toxoplasma gondii* Tissue Cysts. Annu Int Conf IEEE Eng Med Biol Soc 2021, 3509–3513 (2021).

41. B. C. Place, C. A. Troublefield, R. D. Murphy, A. P. Sinai, A. R. Patwardhan, Machine learning based classification of mitochondrial morphologies from fluorescence microscopy images of *Toxoplasma gondii* cysts. PloS one 18, e0280746 (2023).

42. S. W. Perry, J. P. Norman, J. Barbieri, E. B. Brown, H. A. Gelbard, Mitochondrial membrane potential probes and the proton gradient: a practical usage guide. BioTechniques 50, 98–115 (2011).

43. D. Ghosh, J. L. Walton, P. D. Roepe, A. P. Sinai, Autophagy is a cell death mechanism in *Toxoplasma gondii*. Cellular microbiology 14, 589–607 (2012).

44. B. Anderson-White et al., Cytoskeleton assembly in *Toxoplasma gondii* cell division. International review of cell and molecular biology 298, 1–31 (2012).

45. B. R. Anderson-White et al., A family of intermediate filament-like proteins is sequentially assembled into the cytoskeleton of *Toxoplasma gondii*. Cellular microbiology 13, 18–31 (2011).

46. B. Pfister, S. C. Zeeman, Formation of starch in plant cells. Cell Mol Life Sci 73, 2781–2807 (2016).

47. P. J. Roach, A. A. Depaoli-Roach, T. D. Hurley, V. S. Tagliabracci, Glycogen and its metabolism: some new developments and old themes. The Biochemical journal 441, 763–787 (2012).

48. S. Streb, S. C. Zeeman, Starch metabolism in Arabidopsis. The Arabidopsis book / American Society of Plant Biologists 10, e0160 (2012).

49. A. D. Uboldi et al., Regulation of Starch Stores by a Ca(2+)-Dependent Protein Kinase Is Essential for Viable Cyst Development in *Toxoplasma gondii*. Cell host & microbe 18, 670–681 (2015).

50. W. Bohne, J. Heesemann, U. Gross, Reduced replication of *Toxoplasma gondii* is necessary for induction of bradyzoite-specific antigens: a possible role for nitric oxide in triggering stage conversion. Infection and immunity 62, 1761–1767 (1994).

51. A. V. Naumov et al., Restriction Checkpoint Controls Bradyzoite Development in *Toxoplasma gondii*. Microbiol Spectr 10, e0070222 (2022).

52. S. S. Lin, U. Gross, W. Bohne, Type II NADH dehydrogenase inhibitor 1-hydroxy-2-dodecyl-4(1H)quinolone leads to collapse of mitochondrial inner-membrane potential and ATP depletion in *Toxoplasma gondii*. Eukaryotic cell 8, 877–887 (2009).

53. J. Yang, Z. He, C. Chen, J. Zhao, R. Fang, Starch Branching Enzyme 1 Is Important for Amylopectin Synthesis and Cyst Reactivation in *Toxoplasma gondii*. Microbiol Spectr 10, e0189121 (2022).

54. J. Yang et al., *Toxoplasma gondii* alpha-amylase deletion mutant is a promising vaccine against acute and chronic toxoplasmosis. Microb Biotechnol 13, 2057–2069 (2020).

55. J. L. Wang et al., The protein phosphatase 2A holoenzyme is a key regulator of starch metabolism and bradyzoite differentiation in *Toxoplasma gondii*. Nat Commun 13, 7560 (2022).

56. M. Zhao et al., PP2Acalpha-B’/PR61 Holoenzyme of *Toxoplasma gondii* Is Required for the Amylopectin Metabolism and Proliferation of Tachyzoites. Microbiol Spectr 11, e0010423 (2023).

57. C. W. Vander Kooi et al., From the Cover: Structural basis for the glucan phosphatase activity of Starch Excess4. Proceedings of the National Academy of Sciences of the United States of America 107, 15379–15384 (2010).

58. C. A. Worby, M. S. Gentry, J. E. Dixon, Laforin: A dual specificity phosphatase that dephosphorylates complex carbohydrates. J. Biol. Chem. 281, 30412–30418 (2006).

59. Y. Gibon et al., Adjustment of growth, starch turnover, protein content and central metabolism to a decrease of the carbon supply when Arabidopsis is grown in very short photoperiods. Plant Cell Environ 32, 859–874 (2009).

60. A. Pokhilko, A. Flis, R. Sulpice, M. Stitt, O. Ebenhoh, Adjustment of carbon fluxes to light conditions regulates the daily turnover of starch in plants: a computational model. Molecular bioSystems 10.1039/c3mb70459a (2014).

61. A. M. Smith, M. Stitt, Coordination of carbon supply and plant growth. Plant Cell Environ 30, 1126–1149 (2007).

62. S. C. Zeeman, S. M. Smith, A. M. Smith, The diurnal metabolism of leaf starch. The Biochemical journal 401, 13–28 (2007).

63. S. Kato, H. G. Nam, The Cell Division Cycle of Euglena gracilis Indicates That the Level of Circadian Plasticity to the External Light Regime Changes in Prolonged-Stationary Cultures. Plants (Basel) 10 (2021).

64. M. Sasai, M. Yamamoto, Innate, adaptive, and cell-autonomous immunity against *Toxoplasma gondii* infection. Exp Mol Med 51, 1–10 (2019).

65. M. Sasai, M. Yamamoto, Anti-Toxoplasma host defense systems and the parasitic counterdefense mechanisms. Parasitol Int 89, 102593 (2022).

66. I. A. Khan, S. Hwang, M. Moretto, *Toxoplasma gondii*: CD8 T Cells Cry for CD4 Help. Front Cell Infect Microbiol 9, 136 (2019).

67. I. A. Khan, M. Moretto, Immune responses to *Toxoplasma gondii*. Curr Opin Immunol 77, 102226 (2022).

68. J. P. Gigley, R. Bhadra, M. M. Moretto, I. A. Khan, T cell exhaustion in protozoan disease. Trends in parasitology 28, 377–384 (2012).

69. R. Porte et al., Protective function and differentiation cues of brain-resident CD8+ T cells during surveillance of latent *Toxoplasma gondii* infection. Proceedings of the National Academy of Sciences of the United States of America 121, e2403054121 (2024).

70. M. S. Behnke, J. P. Dubey, L. D. Sibley, Genetic Mapping of Pathogenesis Determinants in *Toxoplasma gondii*. Annu Rev Microbiol 70, 63–81 (2016).

71. J. P. Dubey, Comparative infectivity of *Toxoplasma gondii* bradyzoites in rats and mice. The Journal of parasitology 84, 1279–1282 (1998).

72. P. Saraf, E. K. Shwab, J. P. Dubey, C. Su, On the determination of *Toxoplasma gondii* virulence in mice. Experimental parasitology 174, 25–30 (2017).

73. A. P. Sinai, E. S. Suvorova, The RESTRICTION checkpoint: a window of opportunity governing developmental transitions in *Toxoplasma gondii*. Current opinion in microbiology 58, 99–105 (2020).

74. C. J. Beckers et al., Inhibition of cytoplasmic and organellar protein synthesis in *Toxoplasma gondii*. Implications for the target of macrolide antibiotics. J Clin Invest 95, 367–376 (1995).

