## Supplementary figures and images for "Dynamics of amylopectin granule accumulation during the course of the chronic *Toxoplasma* infection is linked to intra-cyst bradyzoite replication"

### Supplemental Figure 1

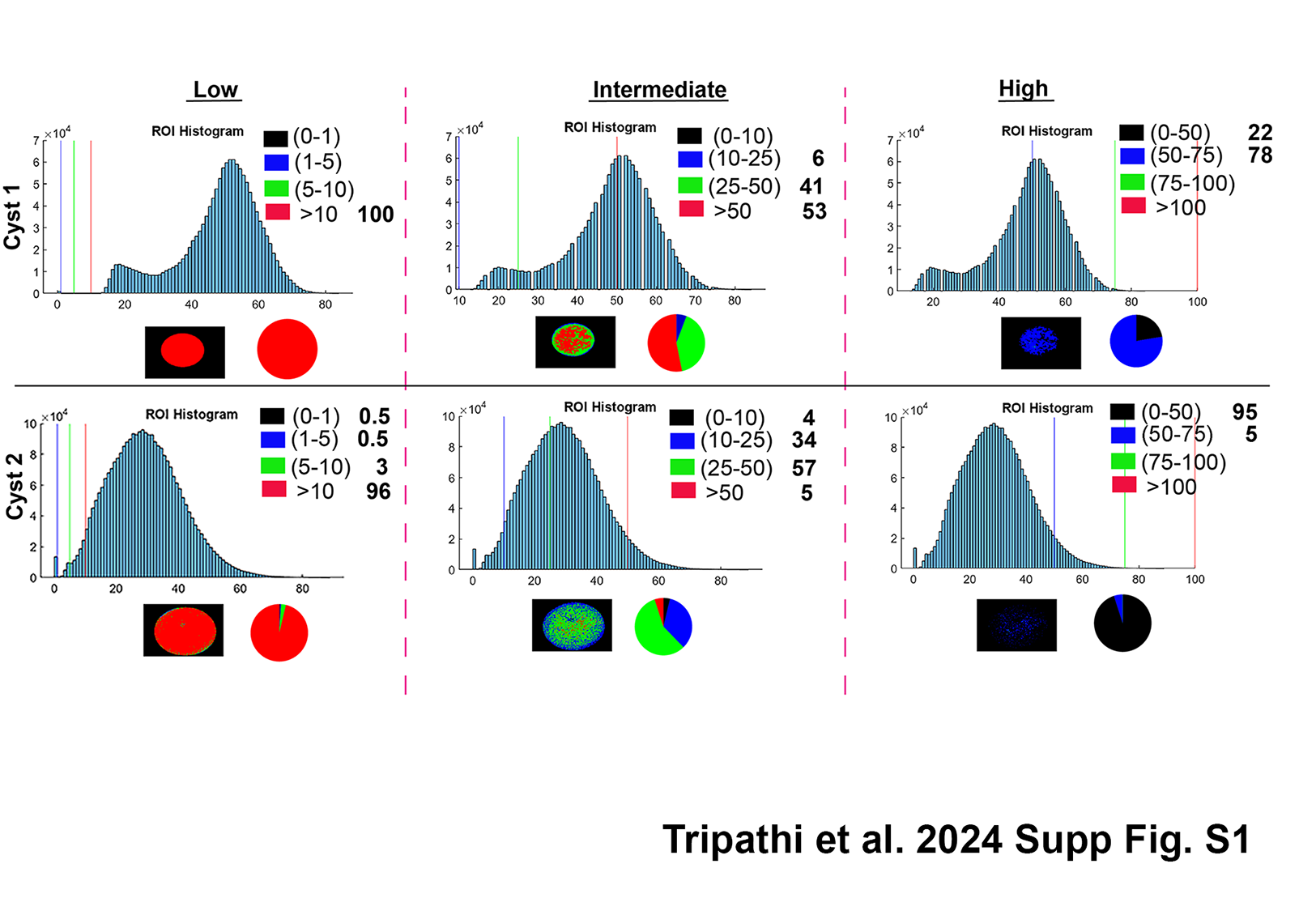

### Supplemental Figure 2

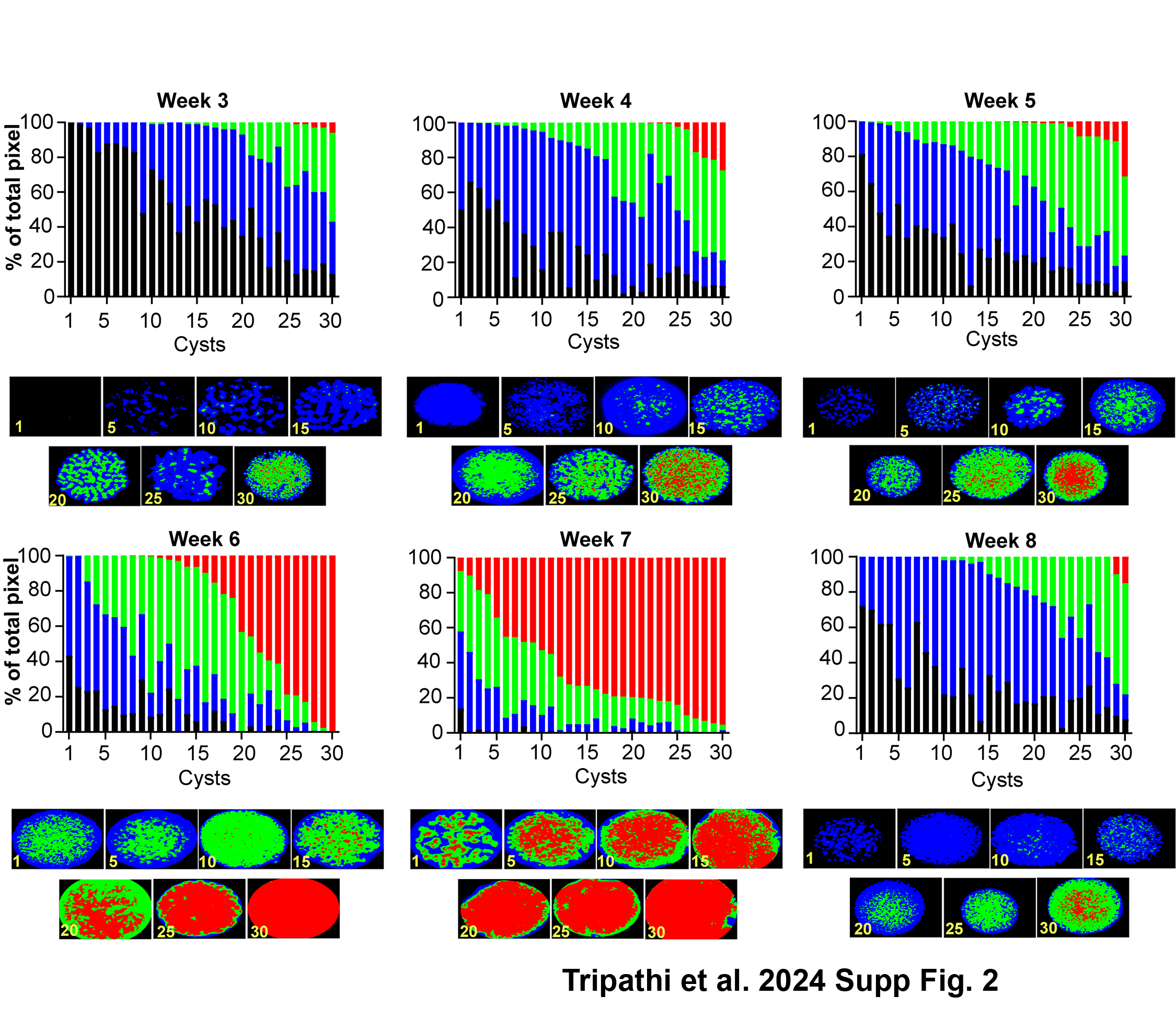

### Supplemental Figure 3

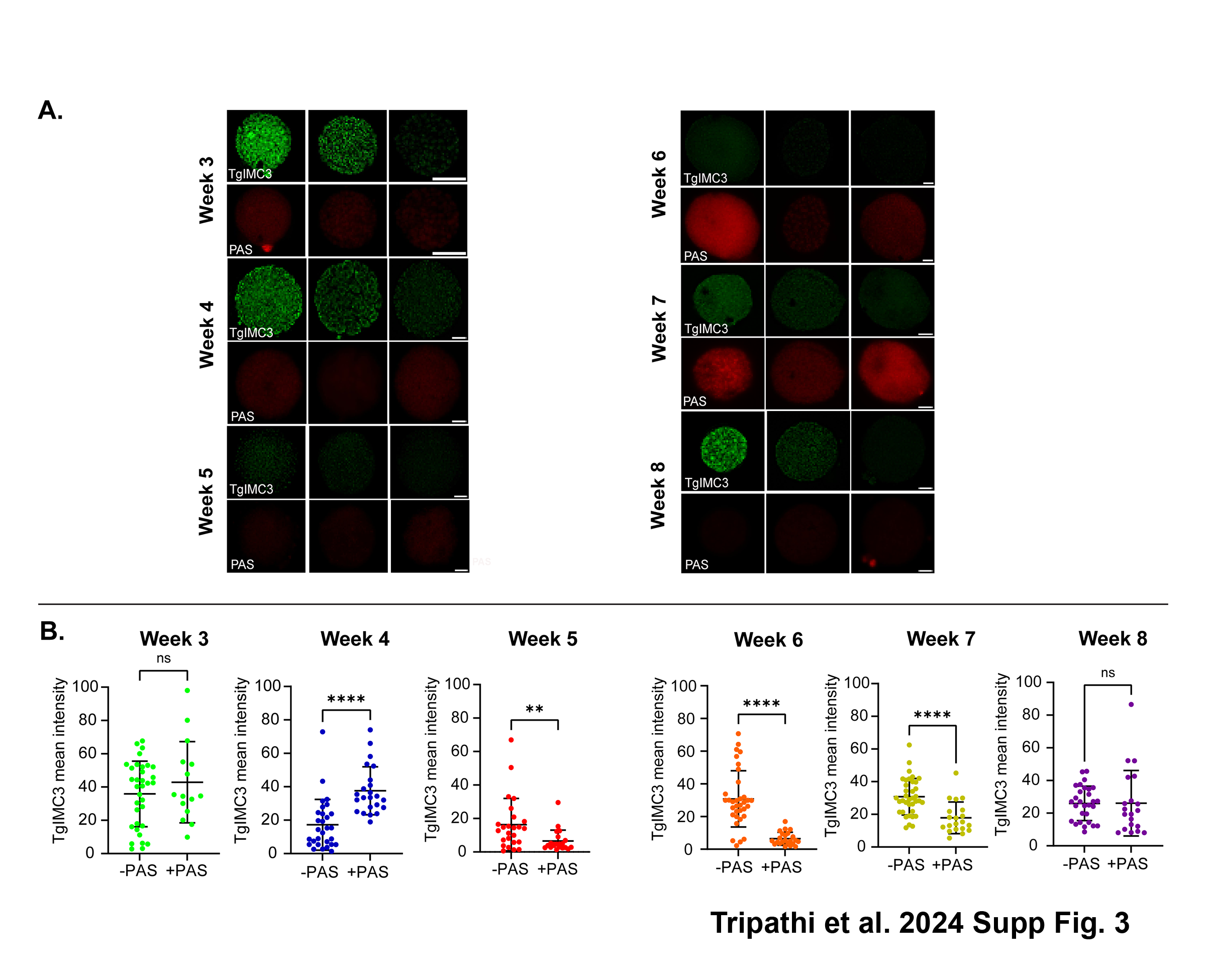

### Supplemental Figure 4

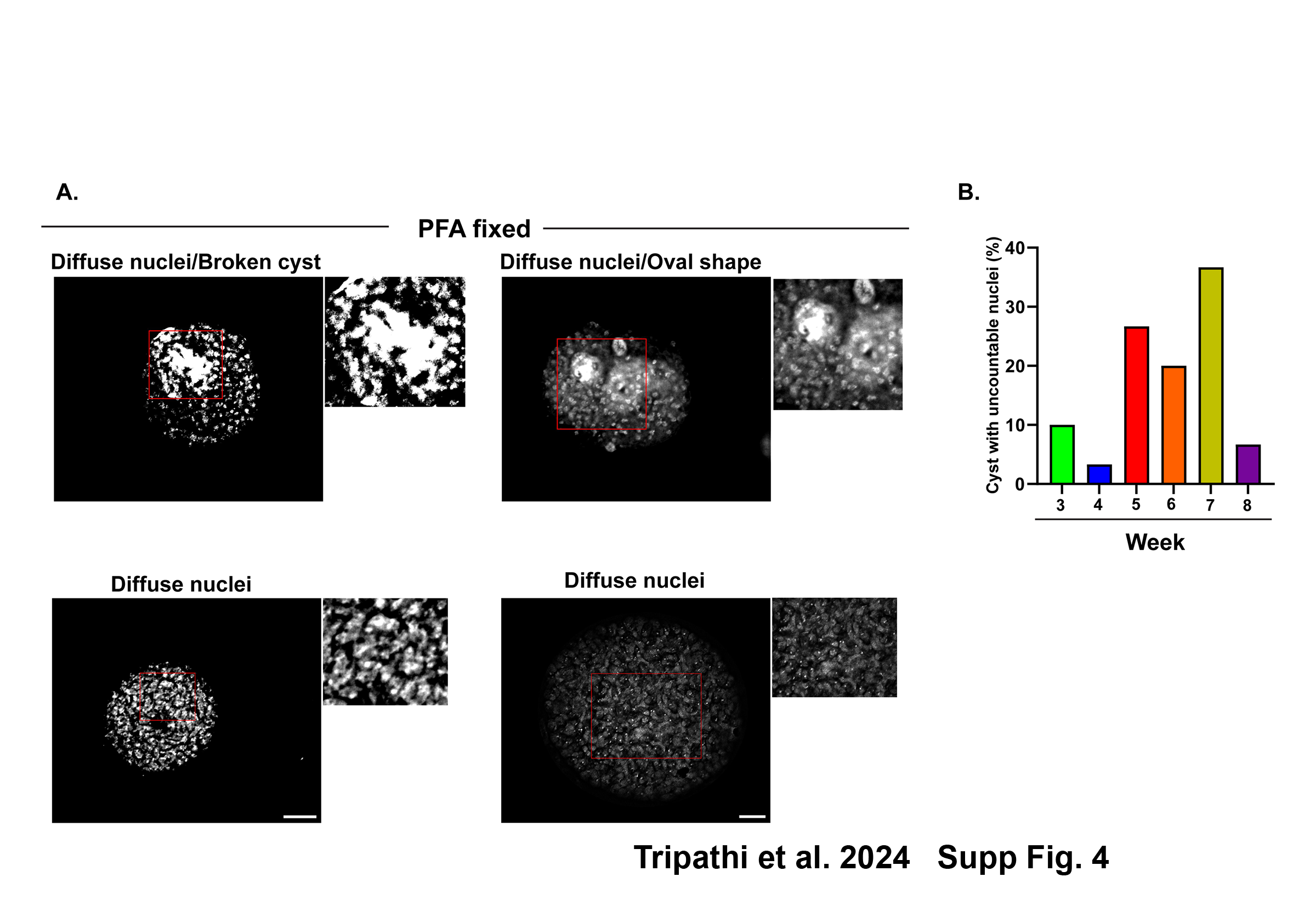
